# SpaGRD deciphers signaling architectures in spatial transcriptomics using graph reaction-diffusion systems

**DOI:** 10.64898/2026.06.28.735031

**Authors:** Jixin Liu, Shuli Sun, Zhitong Chen, Zhengliang Lv, Shuai Jiang, Guojun Li, Bingqiang Liu

## Abstract

The rapid emergence of spatial transcriptomics offers unprecedented opportunities to study cell-cell communication (CCC) by capturing gene expression alongside spatial context. However, existing CCC inference methods often rely on static, heuristic models that overlook the inherently spatiotemporal dynamics and mechanistic complexity of intercellular signaling, limiting both accuracy and biological interpretability. Here, we present SpaGRD, a first-principles-based method that explicitly models ligand-receptor interactions through partial differential equations derived from Fick’s law of diffusion and the mass action law. Leveraging graph signal processing techniques, SpaGRD solves these equations on spatial graphs, providing a principled and generalizable approach to CCC inference. Through extensive simulations, SpaGRD demonstrates superior accuracy and robustness compared to existing methods. Applications to multiple datasets across diverse tissues and platforms reveal dynamic CCC patterns with spatially resolved signaling heterogeneity, providing biologically meaningful insights into cellular coordination and developmental processes. By bridging physical modeling with spatial transcriptomics, SpaGRD provides an accurate, interpretable, and mechanistically grounded framework for advancing quantitative studies of spatiotemporal cell-cell communication.

## Introduction

Multicellular organisms rely on coordinated cellular systems to execute complex biological functions, with such coordination critically dependent on intercellular communication^1^. Such coordination is critically dependent on cells’ capacity for intercellular communication. Cell–cell communication (CCC) is regulated by biochemical signaling, primarily through ligand–receptor binding that triggers downstream responses to guide diverse cellular decisions, playing a crucial role in physiological processes such as homeostasis, embryonic development, and disease progression^2–5^. A widely adopted strategy to describing CCC events involves modeling ligand–receptor interactions between sender and receiver cells^6^. Numerous computational studies on cell–cell communication (CCC) have been conducted based on single-cell RNA sequencing (scRNA-seq) data, such as CellPhoneDB^7^, CellChat^4^, CellCall^8^, SingleCellSignalR^9^, NicheNet^10^. Most of these methods infer CCC based on the assumption that highly co-expressed ligands and receptors are likely to mediate communication^6, 11^. For instance, CellChat infers cell–cell communication by modeling ligand–receptor interactions using mass action models combined with statistical significance testing^4^. Although widely embraced, scRNA-seq-based approaches ignored spatial context, so ligand–receptor pairs can be flagged as interacting even when the corresponding cells are physically separated, resulting in appreciable false-positive rates^6, 12, 13^.

In recent years, the rapid development and large-scale application of spatial transcriptomics (ST) have enabled more accurate descriptions of CCC by preserving valuable spatial information at cell or spot resolution. ST technologies have also greatly accelerated the development of computational methods for CCC inference^14^. Several tools have been proposed from multiple perspectives to integrate gene expression with spatial context, thereby enhancing the precision and reliability of CCC predictions. For instance, SpaTalk models ligand–receptor–target signaling networks between spatially proximal cells using knowledge graphs^11^; SpaOTsc employs an optimal transport framework to infer CCC by linking ligands, receptors, and downstream targets^12^; SpatialDM applies a bivariate Moran’s statistic to detect spatially co-expressed ligand–receptor pairs^15^; COMMOT infers CCC using the spatial expression of ligand and receptor species through a collective optimal transport model^16^; and Spacia leverages a multiple-instance learning framework to model CCC from ST data^17^. These ST-specific methods represent substantial advances in both the prediction and interpretation of cell-cell communication (CCC), achieved by incorporating spatial context into computational models.

Nonetheless, accurately capturing the spatiotemporal dynamics of intercellular signaling to model communication strength remains a significant challenge. Some of the existing methods, such as SpatialDM, incorporate spatial proximity through reciprocal or Gaussian kernel functions that weight pairwise distances, thereby emphasizing interactions between nearby cells. However, these approaches typically use spatial distance as a static weight and fall short of capturing the dynamic influence of the surrounding microenvironment or long-range effects. As a result, communication strength is often treated as fixed, independent of the broader tissue context or potential intercellular competition. Other methods, like COMMOT, employ optimal transport to model CCC from a systems-level perspective, offering a valuable step toward more holistic modeling. Yet, such strategies may impose communication patterns shaped by human-defined assumptions or optimization objectives, potentially diverging from biological reality. Actually, ligand diffusion and molecular interactions in tissues follow physical principles that govern how signals propagate through space, not merely spatial proximity or transport cost. Thus, fully harnessing spatial information in CCC inference requires models that explicitly capture spatial dynamics and physical laws underlying intercellular signaling.

To address these critical gaps, we propose SpaGRD, a first-principles-based framework for inferring cell–cell communication that explicitly models the physical and biochemical processes governing molecular signaling. Unlike methods that rely on heuristic spatial weighting or abstract optimization objectives, SpaGRD constructs a dynamic system of partial differential equations (PDEs) derived from Fick’s law of diffusion and the mass action law to capture how ligands, receptors, and their complexes evolve in space and time. This physically grounded formulation enables a mechanistic understanding of spatial signaling dynamics, accounting for local concentrations, spatial gradients, and microenvironmental influences that simpler models often overlook. To ensure broad applicability across diverse spatial transcriptomics platforms, SpaGRD integrates graph signal processing (GSP) to solve the PDE system directly on spatial graphs, making it compatible with both regular grids (e.g., VisiumHD) and irregular spatial layouts (e.g., Xenium). By unifying physical laws with graph-based computation, SpaGRD offers a general, interpretable, and biologically realistic approach to modeling spatial cell–cell communication.

Building on this mechanistic foundation, SpaGRD generates a dynamic, fine-grained molecular-level representation of spatial signaling, capturing the evolving concentrations and interaction states of ligands, receptors, and their complexes over time. This molecule-centric modeling framework provides a rich and biologically interpretable signal landscape, which can be seamlessly integrated with downstream graph signal processing (GSP)–based analytical modules, such as ligand–receptor module detection or signaling pathway reconstruction. In addition, SpaGRD supports cell-centric communication analysis through a divide-and-conquer strategy that traces the spatial flow of signaling molecules across the tissue graph, enabling the precise identification of active sender and receiver cells. This dual-level perspective—from molecules to cells—allows SpaGRD to bridge mechanistic modeling with interpretable, cell-level communication networks.

Evaluations on simulated spatial datasets reveal that SpaGRD faithfully reconstructs cell–cell communication (CCC) and consistently outperforms existing methods in accuracy. Its accuracy closely matches that of the finite difference method (FDM), a gold-standard numerical technique for solving partial differential equations. In addition, SpaGRD exhibits broad compatibility with diverse spatial transcriptomics platforms, providing a flexible and generalizable solution adaptable to different spatial resolutions and tissue architectures. We applied SpaGRD to a wide range of real-world datasets spanning multiple technologies, including Visium, VisiumHD, Xenium, and Stereo-seq, and biological systems such as brain, tumor, and embryonic tissues. These applications highlight SpaGRD’s ability to capture dynamic, spatially resolved CCC events, leading to biologically meaningful insights into tissue organization, signaling heterogeneity, and developmental processes. By integrating physical laws with graph signal processing, SpaGRD establishes a first-principles-based framework for CCC inference that combines high accuracy, strong interpretability, and robust cross-platform applicability, offering a new paradigm for studying cell–cell communication in spatial omics.

## Results

### Overview of SpaGRD

A key aspect of understanding cell–cell communication is deciphering the interactions between ligands from sender cells and receptors from receiver cells. The input to SpaGRD includes a gene expression matrix (used to approximate ligand and receptor abundance), a spatial coordinates matrix, and prior information on ligand–receptor interactions (**Fig. 1a**). The default ligand–receptor interaction database is CellChatDB^4^, and SpaGRD also supports user-defined ligand–receptor pairs, providing flexibility for customized analyses. First, ligands are produced by sender cells and released into the microenvironment. These ligands then diffuse through the tissue, a process that can be described by Fick’s second law, which explains how concentration changes over time and space. Receptors specifically interact with the diffused ligands, and this binding process can be modeled by the law of mass action (**Fig. 1b-c**). The spatial distribution of receptors influences the diffusion of ligands, as regions with receptor accumulation can locally trap or consume ligands, thereby altering their diffusion dynamics —an aspect that is often overlooked by many cell-cell communication (CCC) inference methods. Together, these complex processes form a dynamic system that can be described by the following PDEs:

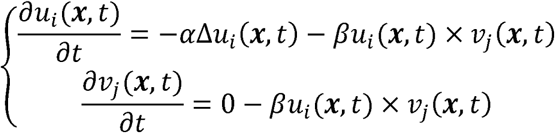

where *u_i_*(***x***, *t*) and *v_j_*(***x***, *t*) represent the concentrations of the ligand *i* and receptor *j* on spatial location *x* at time *t* respectively. Δ*u_i_*(***x***, *t*) is the diffusion term, governed by Fick’s second Law, where Δ is the Laplacian operator, capturing the spatial diffusion process. And *u_i_*(***x***, *t*) × *v_j_*(***x***, *t*) is the reaction term, modeling ligand–receptor interaction based on mass action law. In biological systems, ligand–receptor interactions are often not strictly one-to-one. To account for this, we extend the dynamic system to incorporate more complex interactions, allowing a ligand to interact with multiple receptors and vice versa (see Methods).

**Fig. 1.**
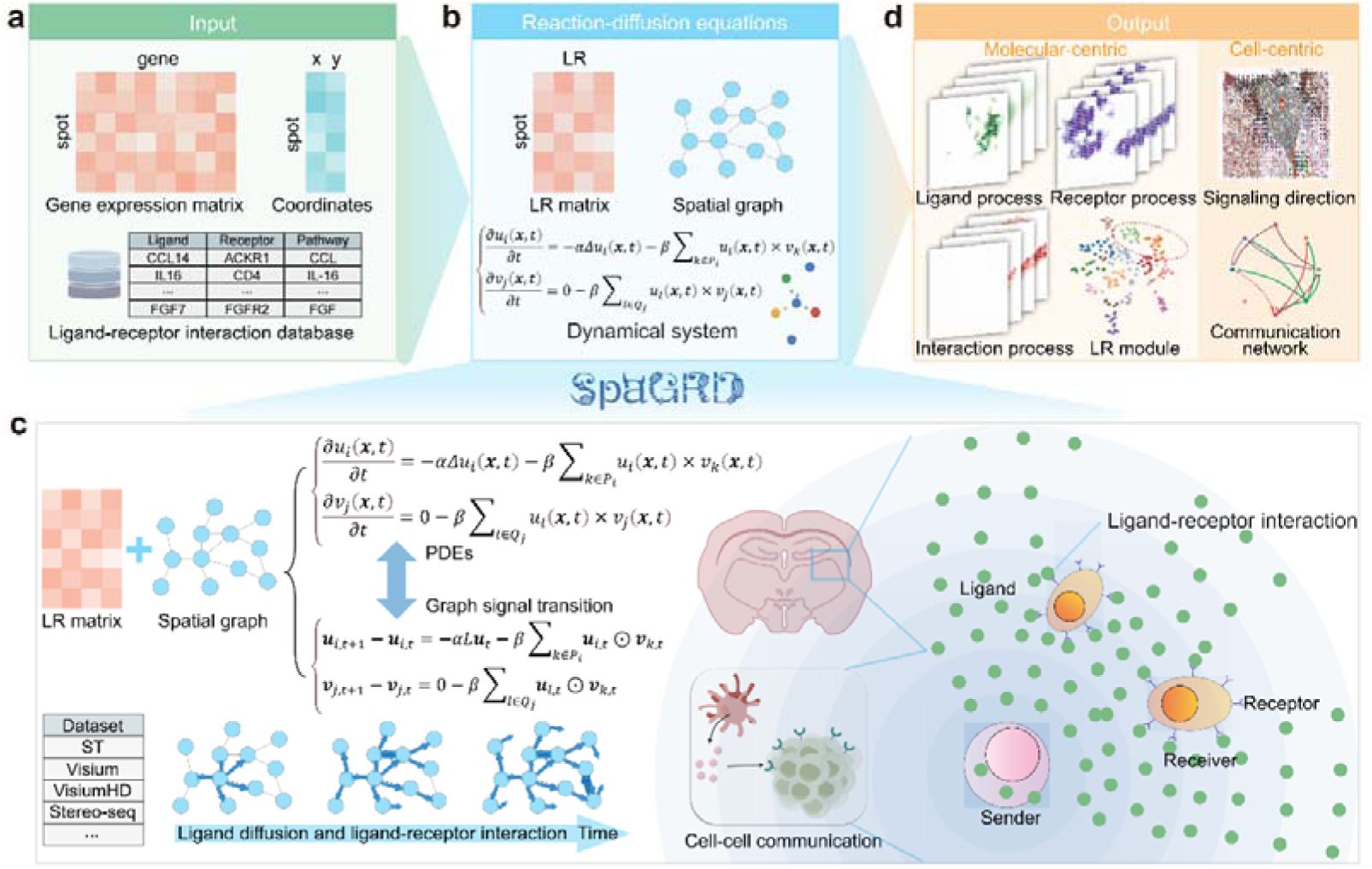
| Overview of the SpaGRD framework. **a**, Input data for SpaGRD includes a curated ligand–receptor interaction database and spatial transcriptomics data, consisting of gene expression matrices and corresponding spatial coordinates. **b**, The reaction–diffusion equations are constructed to infer the spatial dynamics of ligands, receptors and ligand–receptor interactions. SpaGRD formulates a system of partial differential equations (PDEs) based on Fick’s law for ligand diffusion and the law of mass action for ligand–receptor binding. **c**, The system is efficiently solved using graph-based discretization and graph signal transition, enabling direct and scalable solution of the PDEs. The local accumulation of receptors can significantly impede ligand diffusion, as they interact with and sequester diffusing ligands, thereby depleting the concentration of diffusible ligand molecules. **d**, Downstream applications of SpaGRD include pathway-level analysis, interaction module detection, spatial pattern discovery, and cell-type communication inference.

A common mathematical approach to solving these equations is the finite difference method (FDM). Although elegant in form, these equations are not directly applicable to spatial transcriptomics data. One major challenge is that FDM typically relies on mesh generation, whereas the spatial organization of cells or spots varies widely across different technologies. To overcome this, we introduce graph signal processing to solve the equations (**Fig. 1c**). Graphs provide a natural framework to represent spatial relationships between spots or cells, where nodes correspond to cells/spots and edges reflect their spatial neighborhood. In this framework, ligand or receptor levels, approximated by corresponding coding gene expressions, can be treated as graph signals. We thus reformulate the equations into graph-signal-based forms to solve the dynamic system. Such graph signal transitions endow SpaGRD with the flexibility to be applicable to any spatial transcriptomics technology.

Molecular-centric analysis is based on the distribution of ligands, receptors, genes, and their interactions across all cells or spots (**Fig. 1d**). We track ligand–receptor reactions at each time point to capture their dynamic changes. By aggregating reactions across all time points until the system reaches stability, we infer the spatial distribution of ligand–receptor interaction strengths for each ligand–receptor pair at every location. Building on SpaGFT^18^, which utilizes graph Fourier transform for spatial omics representation, these ligand–receptor interaction signals can be analyzed in the frequency domain. For example, spatially variable ligand–receptor pairs can be identified and grouped into modules exhibiting consistent spatial patterns, referred to as LR modules. Together, these analytical approaches offer a powerful molecular-centric framework for studying cell–cell communication events.

Cell-centric analysis is based on ligand flow and ligand–receptor interactions between cells (**Fig. 1d**). Apart from molecular-centric analysis, which describes the spatial distribution of interaction events, another key goal is to identify which individual cells or cell types engage in interactions with one another, achieving cell–cell communication analysis at single-cell resolution. Thanks to the linearity of ligands in PDEs, SpaGRD enables tracing interactions between individual cells via a divide-and-conquer approach. By decomposing a ligand graph signal vector into a set of vectors corresponding to individual cells or spots and solve these equations individually, we can quantify interactions between cell or spot pairs. Since the initial graph signal vector corresponds to a single cell/spot, all subsequent interactions can be interpreted as interactions between that cell/spot and the others. In this way, we can obtain the cell-cell communication matrix at single cell/spot level. Drawing on a similar procedure used in COMMOT^16^, the spatial signaling direction is then obtained by interpolating the CCC matrix into a vector field to identify directionality. Further, by summarizing or averaging cell–cell interaction scores according to cell or spot labels and applying permutation tests, SpaGRD provides a depiction of communication network at the cell-type level. Detailed methods are provided in the Methods section.

### Benchmarking results highlight the superior accuracy and efficiency of SpaGRD compared to existing cell–cell communication inference tools

We evaluated the accuracy of SpaGRD in comparison with existing methods by contrasting the predicted ligand–receptor interaction intensity against the standard numerical solutions of PDEs, following a similar evaluation strategy as used in COMMOT. Three competitive methods, COMMOT, Spacia, and SpatialDM, were included in the comparison, as they are capable of providing cell-resolution descriptions of ligand–receptor interactions. To begin with, we designed ten simulation scenarios involving varying ligand/receptor abundances and interaction rules to test the flexibility of the methods (**Fig. 2a**). For each scenario, ten replicates were generated to further assess robustness. The initial ligand/receptor distributions were simulated using simstpy^19^, a tool designed for spatial omics data simulation. The standard finite difference method (FDM) was then applied to compute the ligand–receptor interaction intensity distributions across individual cells to obtain the numerical solutions, which were designated as the ground truth. Two evaluation metrics, Pearson correlation coefficient (PCC) and root mean squared error (RMSE), were subsequently used to quantify the consistency of predicted interaction intensities with the ground truth for each ligand–receptor pair.

**Fig. 2.**
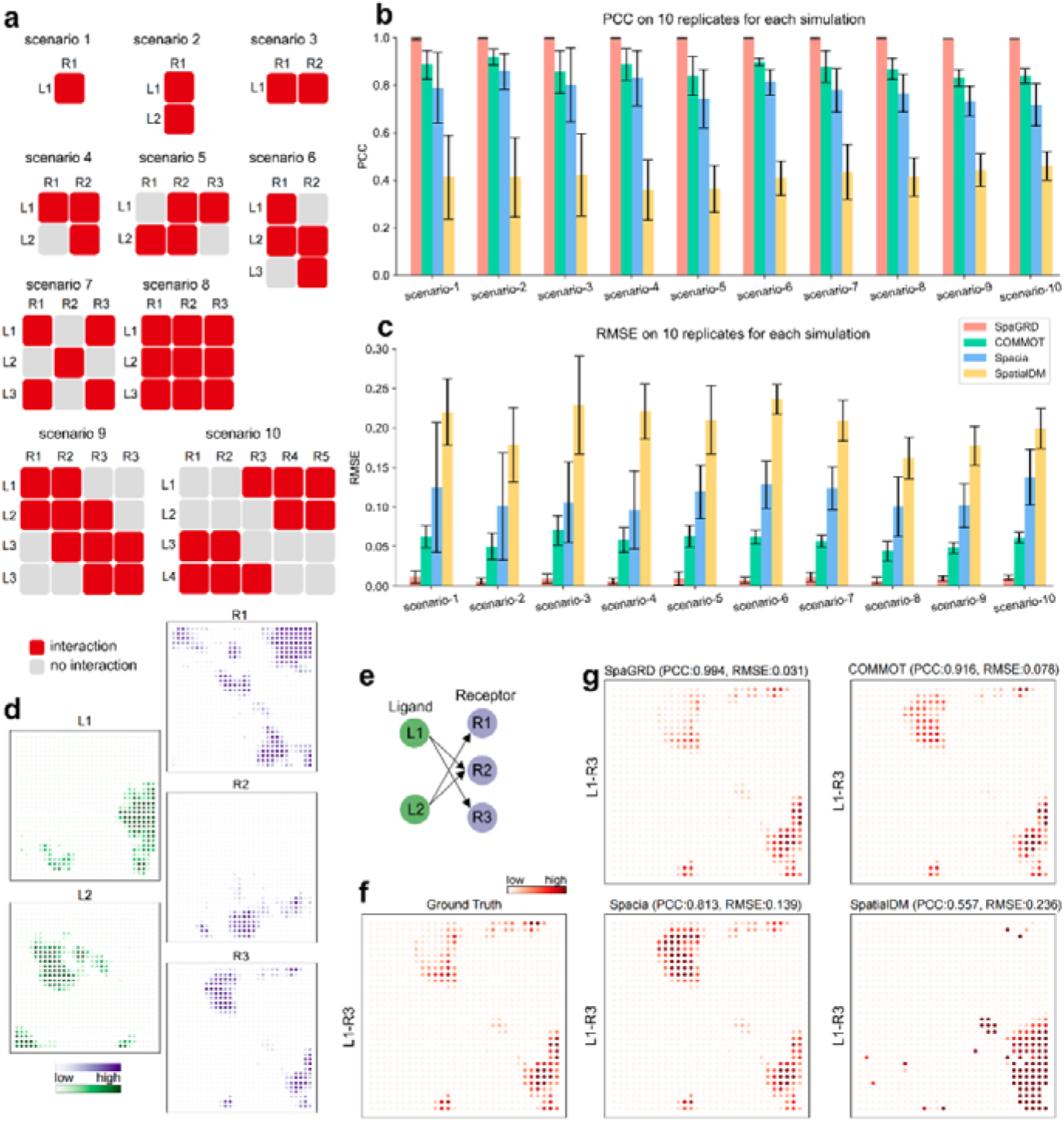
| Benchmarking results showed superior performance against the existing methods. **a**, Ten simulation scenarios were designed to cover a diverse range of ligand–receptor interaction scenarios. Here, ‘L*i*’ denotes ligand *i* and ‘R*j*’ denotes receptor *j*. A red connection between ligand *i* and receptor *j* indicates that they are capable of interacting with each other. **b**, The Pearson Correlation Coefficient (PCC) was used to evaluate performance. Ten replicates were generated for each simulation scenario. For each ligand–receptor pair, evaluation metrics were calculated by comparing the predicted interaction intensities with the ground truth, derived from numerical solutions of partial differential equations (PDEs). **c**, Performance was evaluated using the Root Mean Square Error (RMSE). Ten replicates were generated for each simulation case. To enable fair comparisons across methods, predicted interaction intensities were normalized so that their maximum values aligned with those of the ground truth. **d**, An example of the spatial distributions of ligand and receptor expressions in a simulated dataset from simulation scenario 5. **e**, Schematic diagram of ligand–receptor interactions. L1 and L2 represent two ligands, and R1–R3 represent three receptors. Arrows indicate potential interactions between ligands and receptors, demonstrating a many-to-many interaction scenario. **f,** Ground truth interaction intensity between L1 and R3, obtained by numerical simulation of PDEs. **g,** Predicted interaction intensities between L1 and R3 by four methods: SpaGRD, COMMOT, Spacia, and SpatialDM. Each panel includes the Pearson correlation coefficient (PCC) and root mean square error (RMSE) with respect to the ground truth. Color intensity indicates the predicted strength of interaction at each spatial location.

The results demonstrated that SpaGRD significantly outperforms existing methods, achieving a leading edge in performance (**Fig. 2b–c**), primarily due to its first-principle-based modeling. The PCC values for nearly all ligand–receptor interaction pairs are close to 1, indicating strong agreement with the ground truth. Pairwise comparisons between SpaGRD and other methods further highlight its advantage (**Supplementary Fig. 1a–c** and **Supplementary Fig. 2a–c**). SpatialDM underperforms in this task, possibly due to its statistical framework, which may be better suited for significance testing rather than fine-grained interaction reconstruction. Benchmark results also confirm that the graph-based approach yields interaction patterns highly consistent with those derived from the traditional mesh-grid-based method. As an illustrative example, a replicate from simulation scenario 5 consists of two ligands (L1 and L2) and three receptors (R1, R2, and R3), where L1 interacts with R2 and R3, and L2 with R1 and R2 (**Fig. 2d–e**). For instance, comparing the ground truth interaction distribution of L1–R3 with the predictions from four methods (**Fig. 2f**), SpaGRD achieves the highest PCC (0.994) and the lowest RMSE (0.031). In contrast, COMMOT achieves PCC = 0.916 and RMSE = 0.078, Spacia reaches PCC = 0.813 and RMSE = 0.139, while SpatialDM performs the worst with PCC = 0.557 and RMSE = 0.236 (**Fig. 2g**). The results for other interaction pairs follow a similar trend (**Supplementary Fig. 3a–c**).

In summary, SpaGRD provides highly accurate and efficient characterization of cell–cell interaction events, surpassing existing approaches in both performance and robustness. Notably, SpaGRD maintains extremely high accuracy even while incorporating graph signal transition, which adds flexibility and adaptability across diverse spatial transcriptomics platforms.

### SpaGRD uncovers spatially resolved ligand–receptor interactions and functional heterogeneity in the adult mouse brain

We first applied SpaGRD to an adult mouse brain Visium dataset. The tissue section encompasses multiple anatomical structures, including the cortex, hippocampal formation, thalamus, hypothalamus, and fiber tracts (**Fig. 3a–b**). After gene filtering, 2,702 ligand–receptor pairs were retained for analysis. SpaGRD enables the visualization of individual ligand–receptor interactions at single-cell or spot resolution. For example, we observed strong PSAP–GPR37 interactions in the fiber tracts, suggesting a potential functional role of this signaling axis in the region (**Fig. 3c**). Previous studies have demonstrated the critical role of PSAP–GPR37 signaling in oligodendrocyte lineage cells^20, 21^, a major cell type within fiber tracts. Similarly, high interaction enrichment of NTS–NTSR2 in the hypothalamus is consistent with earlier findings^22, 23^. Notably, PSAP–GPR37 and NTS–NTSR2 display distinct spatial interaction patterns: the former closely aligns with the ligand distribution, while the latter shows a stronger correspondence with receptor localization. Additionally, interactions such as ADCYAP1–ADCYAP1 and FGF–FGFR1 also exhibit characteristic spatial distributions (**Supplementary Fig. 4a**).

**Fig. 3.**
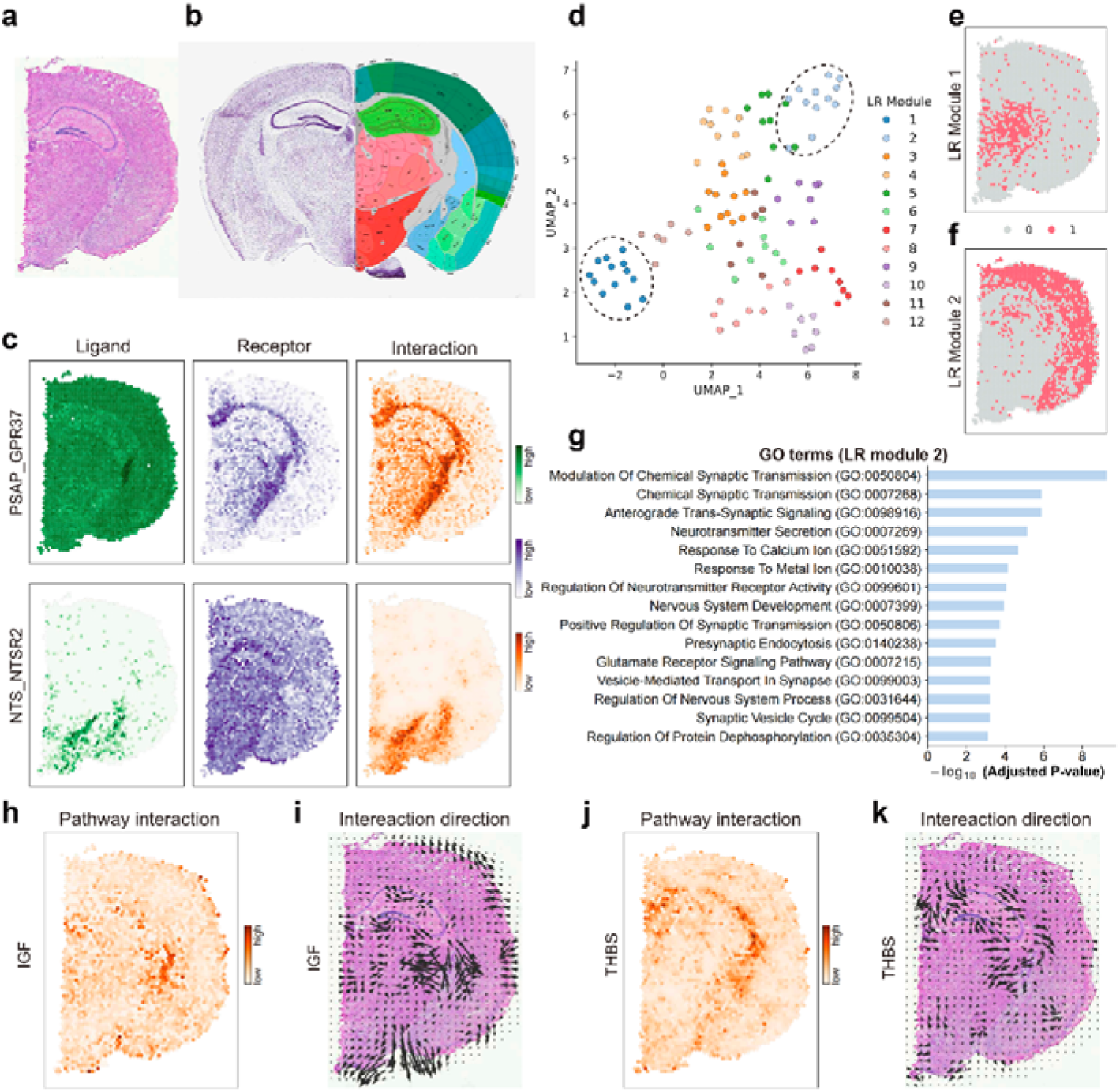
| SpaGRD uncovers spatially resolved ligand–receptor interactions and functional heterogeneity in the adult mouse brain. **a**, Matched histology image of the 10x Visium mouse brain spatial transcriptomics data. **b**, Anatomical annotations from the Allen Mouse Brain Atlas. **c,** Representative spatial distributions of ligands, receptors, and their interactions. For the *PSAP_GPR37* pair, where the ligand gene is *Psap* and the receptor gene is *Gpr37*, the spatial pattern of the interaction closely resembled that of the receptor gene *Gpr37*. In contrast, the spatial distribution of the *NTS_NTSR2* interaction, involving the ligand gene *Nts* and receptor gene *Ntsr2*, aligned more closely with the ligand gene *Nts*. **d,** The UMAP visualization of clustering spatially variable ligand-receptor interactions. 12 ligand-receptor (LR) modules were identified. Each point represents a ligand-receptor pair. **e,** Spots with high interaction intensity for ligand-receptor interactions in LR module 1. **f**, Spots with high interaction intensity for ligand-receptor interactions in LR module 2. **g**, Enriched GO terms of genes positively correlated with ligand-receptor pairs in LR module 2. **h**, Spatial distribution of IGF pathway interactions, calculated by summarizing the interaction intensities of all ligand-receptor pairs involved in the IGF pathway. **i**, Vector fields of the IGF pathway. Arrows indicate the specific direction and intensity of the interactions. **j**, Spatial distribution of THBS pathway interactions, calculated by summarizing the interaction intensities of all ligand-receptor pairs involved in the THBS pathway. **k**. Vector fields of the THBS pathway.

Leveraging the flexibility of graph signal representations, we next explored the spatial patterns of ligand–receptor (LR) interactions on the graph. Using SpaGFT, 106 spatially variable LR interaction pairs were identified. By clustering these interactions based on their frequency-domain representations, 12 distinct LR modules were detected (**Fig. 3d**). Module 1, which includes representative interactions such as NRTN–GFRA2, WNT3–FZD3–LRP6, and WNT4–FZD10–LRP6, is predominantly located in the thalamus (**Fig. 3e** and **Supplementary Fig. 4b**). In contrast, Module 2, with representative pairs like AGT–MAS1, CCK–CCKBR, and LAMC2–DAG1, is primarily enriched in the cortex (**Fig. 3f** and **Supplementary Fig. 4c**). The formation of these modules suggests that the associated LR pairs may act synergistically within specific brain regions to support regional functions. To further explore their biological relevance, we conducted correlation analysis between LR modules with gene expressions (See “Methods”). For instances, Module 1 showed strong spatial correlation with genes such as *Rgs16*, *Lef1*, and *Zic1* (**Supplementary Fig. 4d**), while Module 2 correlated with *Arpp21*, *Chn1*, and *Mef2c* (**Supplementary Fig. 4e**). To elucidate the functional roles of these LR modules, we performed Gene Ontology (GO) enrichment analysis on the genes correlated with each module. The results revealed that genes associated with Module 1 were enriched in processes related to neuronal development and electrophysiological regulation, including axonogenesis, regulation of cell migration, canonical Wnt signaling, and transmembrane transport of inorganic cations (**Supplementary Fig. 4f**). In contrast, genes correlated with Module 2 were enriched in cell–cell communication (CCC)-related processes, such as the synaptic vesicle cycle, glutamate receptor signaling, and modulation of chemical synaptic transmission (**Fig. 3g**).

As a dynamic system model, SpaGRD is also capable of capturing the dynamic changes in ligand–receptor interactions over time (**Supplementary Fig. 5a**). These dynamic changes in interaction intensity reflect the reaction speed, which may be influenced by interaction distance—with long-range interactions typically requiring more time to stabilize, whereas short-range interactions tend to reach stability more rapidly. To examine whether different ligand–receptor pairs exhibit distinct rates of interaction intensity increase, we divided the first 100 simulation steps into three intervals: T1 (steps 0–5), T2 (steps 6–30), and T3 (steps 31–100), and aggregated the interaction intensity within each period. The results revealed clear differences: some ligand–receptor pairs displayed strong interaction intensity from the outset, while others showed a gradual increase in strength following the initial phase (**Supplementary Fig. 5b**). These findings suggest that ligand–receptor interactions follow diverse dynamic trajectories, potentially reflecting differences in their underlying biological mechanisms.

Lastly, by aggregating ligand–receptor interactions associated with the same pathway, pathway-level interaction intensity profiles were generated. SpaGRD effectively captured the complex spatial distribution patterns and interaction directions of key signaling pathways. For example, the IGF pathway (**Fig. 3h–i**), which is known to promote the survival and proliferation of neural cells and regulate their growth and maturation^24^, exhibited distinct spatial characteristics. Similarly, the THBS pathway (**Fig. 3j–k**), which facilitates synaptogenesis in the central nervous system^25^, also showed unique spatial interaction features.

Overall, SpaGRD enables comprehensive inference of ligand–receptor interaction properties in the mouse brain, supporting in-depth analysis of biological heterogeneity and underlying molecular mechanisms relevant to nervous system function and development.

### SpaGRD reveals CD44-mediated cell–cell communication and intratumor heterogeneity in brain metastasis

In the following, we applied SpaGRD to a brain metastasis Xenium dataset to dissect cell–cell communication within the tumor microenvironment. In this analysis, we focused primarily on tumor-related CCC and manually selected the tumor region and its surrounding areas, encompassing 12,950 cells and 243 genes (**Fig. 4a** and **Supplementary Fig. 6a**). Due to the limited gene coverage of Xenium technology we concentrated on interactions involving tumor cells, identified by the marker gene *Cd44*. As a first step, we examined two representative interactions: COL1A1–CD44 and COL6A1–CD44. The spatial distributions of the ligands and receptors are shown in (**Fig. 4b**). Notably, *Col1a1* was predominantly expressed within the tumor region, especially around fibroblasts. In contrast, *Col6a1* exhibited a distinct spatial pattern, being primarily located outside the tumor area.

**Fig. 4.**
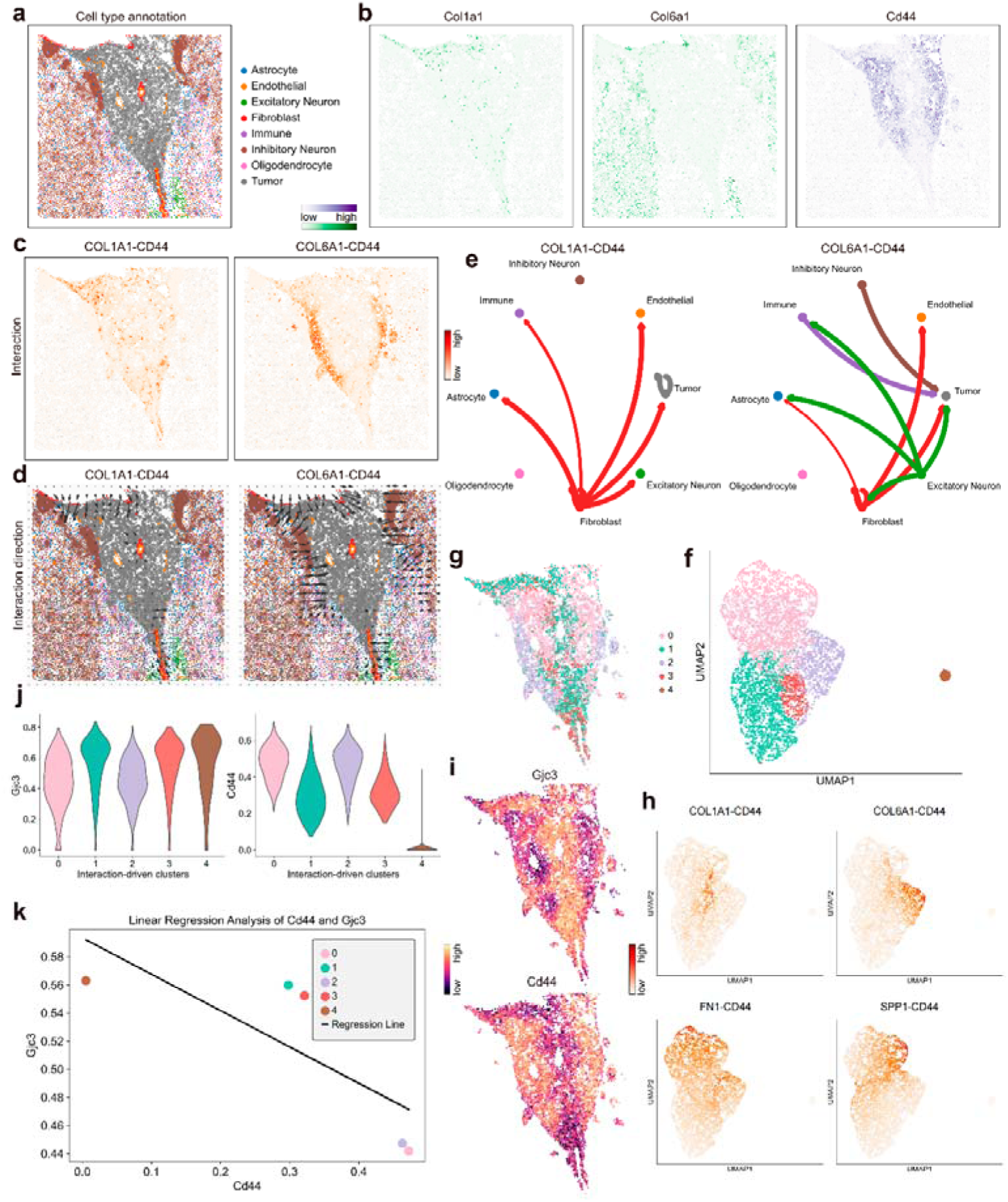
| SpaGRD reveals CD44-mediated cell-cell communication and intratumor heterogeneity in brain metastasis. **a**, Cell type annotation of the selected region in spatial transcriptomics tissue section **b**, Spatial expression patterns of *Col1a1*, *Col6a1*, and *Cd44* genes. **c**, Interaction intensity maps inferred by SpaGRD for the ligand-receptor pairs COL1A1-CD44 and COL6A1-CD44. **d**, Vector fields showing the direction and magnitude of interactions for COL1A1-CD44 (left) and COL6A1-CD44 (right). **e**, Cell-cell communication networks derived from SpaGRD for the COL1A1-CD44 (left) and COL6A1-CD44 (right) pairs. Edge thickness represents interaction intensity, and the arrow direction reflects inferred ligand-to-receptor interactions. **f**, UMAP embedding of spatial spots based on interaction-driven gene expression features, identifying five interaction-driven clusters. **g**, Spatial distributions of the five interaction-driven clusters. **h**, Spatial projection of SpaGRD-inferred interaction strength for additional CD44-related ligand-receptor pairs (FN1-CD44 and SPP1-CD44) in UMAP space. **i**, Spatial expression patterns of *Gjc3* and *Cd44*. **j**, Violin plots showing the expression levels of *Gjc3* (left) and *Cd44* (right) across the five interaction-driven clusters. **k**, Linear regression analysis reveals a negative correlation between *Cd44* and *Gjc3* expression levels across spatial domains.

We applied SpaGRD to investigate how variations in the spatial distributions of ligands may differentially influence their receptor interactions. The results revealed that the COL1A1–CD44 interaction occurred at multiple sites within the tumor, whereas the COL6A1–CD44 interaction showed significantly higher intensity at the tumor margin, indicating a more distinct and localized spatial pattern (**Fig. 4c**). Furthermore, analysis of the interaction dynamics indicated that COL6A1–CD44 required a longer time to reach stability, suggesting a greater interaction distance (**Supplementary Fig. 6b–c**). Interaction directionality analysis showed that COL1A1–CD44 exhibited directional signaling within the tumor, while COL6A1–CD44 displayed clear directional interactions originating from the tumor exterior toward the tumor margin (**Fig. 4d**). This pattern suggests a potential role for COL6A1–CD44 in tumor progression and microenvironment remodeling. This interpretation is consistent with previous studies that have highlighted its involvement in modulating the tumor microenvironment and influencing disease progression^26–28^. Finally, cell type communication network analysis revealed distinct cellular contexts for these interactions (**Fig. 4e**): fibroblasts were the primary sender cells in the COL1A1–CD44 interaction, whereas both fibroblasts and excited neurons contributed significantly to the COL6A1–CD44 interaction.

Next, we explored whether differences in cell–cell interactions contribute to intratumor heterogeneity. In addition to the previously identified interactions, we examined two additional ligand–receptor pairs involving CD44 as the receptor: FN1–CD44 and SPP1–CD44. These interactions exhibited distinct molecular distribution patterns, directionalities, and communication networks, each with unique spatial characteristics (**Supplementary Fig. 7a–e**). Notably, endothelial cells were prominently involved in the FN1–CD44 interaction, consistent with previous findings^29^. To further characterize heterogeneity, we performed clustering analysis based on the interaction profiles of four ligand–receptor pairs (COL1A1–CD44, COL6A1–CD44, FN1–CD44, and SPP1–CD44), identifying five interaction-driven tumor cell clusters (**Fig. 4f**). These clusters displayed spatial organization within the tissue (**Fig. 4g**) and significant differences in their interaction profiles (**Fig. 4h** and **Supplementary Fig. 8a**), as well as distinct spatial patterns (**Fig. 4i**). For example, Cluster 1 exhibited high levels of SPP1–CD44 and FN1–CD44 interactions, whereas Cluster 2 showed elevated COL6A1–CD44 interaction. These variations underscore the complexity of intratumor heterogeneity shaped by cell–cell communication.

Further differential gene expression (DEG) analysis of the interaction-driven clusters revealed distinct transcriptional profiles among tumor cell subpopulations (**Supplementary Fig. 8b**). Interestingly, two DEGs, *Gjc3* and *Cd44*, displayed distinct spatial expression patterns (Fig. 4g): Clusters 1, 3, and 4 showed high expression of *Gjc3*, whereas Clusters 0 and 2 exhibited elevated expression of Cd44 (**Fig. 4j**). Moreover, linear regression analysis based on the interaction-driven clusters revealed a negative correlation between *Cd44* and *Gjc3* expression (**Fig. 4k**), suggesting potential regulatory interplay between these genes in defining tumor subtypes.

In summary, these findings validate the utility of SpaGRD in uncovering tumor heterogeneity and deciphering the tumor microenvironment. By resolving spatially distinct ligand–receptor interactions and communication patterns, SpaGRD reveals functional differences among tumor subpopulations and offers insights into tumor progression and cellular organization.

### SpaGRD deciphers tumor heterogeneity and tumor-border interactions through signal pathway correlation

Cell-cell communication is essential in regulating tumorigenesis, phenotypic maintenance, tumor heterogeneity, and the tumor microenvironment^30^. It mediates communication between cancer cells and surrounding stromal or immune cells, influencing cancer progression, immune evasion, and therapeutic response^31^. The gradient interaction distribution of ligand–receptor pairs in the tumor microenvironment exhibits typical characteristics during the progression of colorectal cancer. In the following, we applied SpaGRD to two human colorectal cancer datasets, including a Visium dataset and a VisiumHD dataset.

In the Visium data, we first identified spatial domains based on gene expression information, where domains 0, 2, 4, 8, 9, and 10 were tumor regions (**Supplementary Fig. 9a**). SpaGRD identified TGFβ (Transforming Growth Factor-beta) and HH (Hedgehog) signal pathways presenting opposite directions. The TGFβ pathway, a core signaling network regulating cell proliferation, differentiation, apoptosis, and microenvironmental interactions, shows a spatial action direction diffusing from stromal cells (such as CAFs) to the tumor interior. Conversely, the spatial action direction of the HH pathway exhibits directional transmission from tumor cells to stromal cells. It activates stromal cells such as CAFs, forming a pro-cancerous cascade reaction (**Fig. 5a–b**). Additionally, SpaGRD identified other signal pathways with opposite directions and spatial characteristics, indicating their association with tumor spatial heterogeneity (**Supplementary Fig. 9b**).

**Fig. 5.**
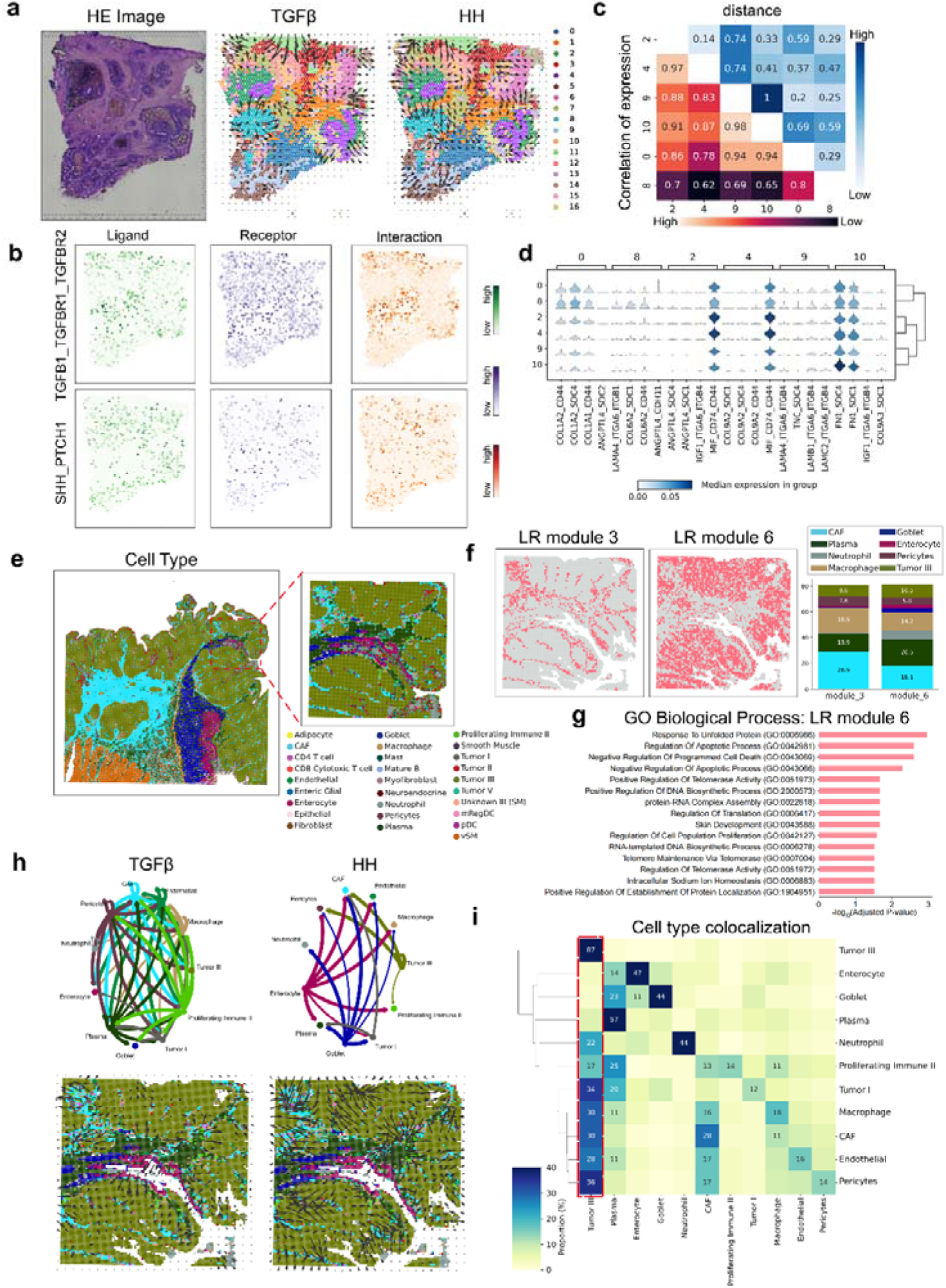
| SpaGRD deciphers tumor heterogeneity and tumor-border interactions. **a**, HE image, and Vector fileds of TGFβ (Transforming Growth Factor-beta) signaling and HH (Hedgehog) signaling, with directed edges representing pathway signaling on cluster map by Leiden in human colorectal cancer Visium data. **b**, Representative spatial distributions of ligand, receptor, and their interactions. For the TGFB1-TGFBR1-TGFBR2 pair, where the ligand gene is *TGFB1*, and the receptor gene is the minimum of and *TGFBR1* and *TGFBR2*, the spatial pattern of the interaction closely resembled that of the receptor gene, the minimum of and *TGFBR1* and *TGFBR2*. In contrast, the spatial distribution of the SHH-PTCH1 interaction, involving the ligand gene *SHH* and receptor gene *PTCH1*, aligned more closely with the ligand gene *SHH*. **c**, The heatmap of the correlation of gene expression and spatial distances among the six tumor subregions. **d**, Stacked violin plots showing the different pathway signal levels across the six tumor subregions. **e**, Cell type annotation in human colorectal cancer VisiumHD data with 16um resolution. The left is original dataset, and the right is selected small region. **f**, Spots with high interaction intensity for ligand-receptor interactions in LR module 3 and 6. Cell type proportion of LR module 3 and 6. **g**, Enriched GO terms of genes positively correlated with ligand-receptor pairs in LR module 6. **h**, Cell-cell communication networks derived from SpaGRD for the TGFβ (left) and HH (right) pathway. Edge thickness represents interaction intensity, and the direction reflects inferred ligand-to-receptor signaling. Vector fields showing the direction and magnitude of interactions for TGFβ (left) and HH (right). **i**, Colocalization heatmap of cell types corresponding to the **h**.

SpaGRD also reveals that both expression similarity and spatial distance in tumor regions influence tumor heterogeneity. The expression and spatial heterogeneity among the six tumor subregions are related to the heterogeneity of signal pathways (**Fig. 5c–d**). Tumors 2 and 4 are the most similar expression and exhibit a chimeric neighboring structure. The MIF-CD74-CD44 interaction, which regulates immune cells and cytokines, shows strong interactions between these two tumors. However, tumors 9 and 10, which are expression-similar but spatially distant, also have similar interaction signals, consistent with the distribution pattern in the image. This suggests the cross-spatial regulatory mechanism and spatial functional coordination strategy of the tumor microenvironment. In contrast, tumors 0 and 8, which are spatially close, exhibit spatial isolation and local specificity.

In the VisiumHD data with near single-cell resolution (16 μm), SpaGRD clearly maps the spatial patterns of tumors and boundaries, as well as tumor-border interactions, through ligand–receptor sets. We selected a subregion with multiple cell types and clustered ligand–receptor pairs to obtain LR modules (**Fig. 5e**). We selected LR module 3 and LR module 6 with similar signaling characteristics as examples (**Supplementary Fig. 9c**), which show significant differences in spatial distribution, cell types, and marked signal pathways (**Fig. 5f** and **Supplementary Fig. 9d**). Further differentiation shows that LR module 6 is associated with the Regulation of Apoptotic Process, while LR module 3 is related to Collagen Fibril Organization and Regulation of Cell Migration, indicating that they characterize the boundary and tumor regions, respectively (**Fig. 5g and Supplementary Fig. 9e**). Additionally, we explored the relationship between cell-type interactions and co-localization **(Fig. 5h–i**). The TGFβ signal pathway shows interactions among almost all listed cell types, while the HH signal pathway shows specificity in interaction with Tumor III. Tumor III has almost no interactions with Goblet, Enterocyte, Plasma, Proliferating immune Il, and Neutrophil, which is positively correlated with cell-type co-localization and avoidance.

### SpaGRD depicts spatiotemporal dynamics of pathway-mediated cell–cell communication during mouse embryogenesis

We applied SpaGRD to investigate ligand–receptor interactions in developmental biology using four Stereo-seq mouse embryo datasets spanning embryonic days E9.5 to E11.5. Spot-level annotations were obtained from the original study^32^ (**Fig. 6a**). SpaGRD effectively inferred ligand–receptor interaction patterns across developmental time points. For instance, IGF2–IGF2R interaction, which can promote cell proliferation and differentiation to regulate embryonic growth^33^, showed certain spatial patterns in ligand and receptor distributions (**Supplementary Fig. 10a–b**). The interaction distributions results showed that it exhibited high interaction intensity in multiple regions, especially in heart (**Fig. 6b**). These findings are consistent with prior studies confirming the regulatory role of IGF2–IGF2R signaling in cardiomyocyte proliferation and heart morphogenesis during development^34–37^. Another notable interaction is SHH–PTCH1, a core signaling axis in vertebrate embryogenesis that guides processes such as craniofacial morphogenesis and gastrointestinal (GI) development^38–42^. Interaction analysis based on ligand gene Shh and receptor gene Ptch1 (**Supplementary Fig. 10d–e**) revealed spatial patterns aligned with these biological roles. Specifically, we observed region-specific interaction intensities in parts of the brain and GI tract (**Fig. 6c** and **Fig. 6f–g**), which are consistent with prior biological insights. Additionally, SpaGRD enables the capture of dynamic interaction changes within single datasets, offering fine-grained insights into ligand–receptor communication over time and space (**Fig. 6h–i**).

**Fig 6.**
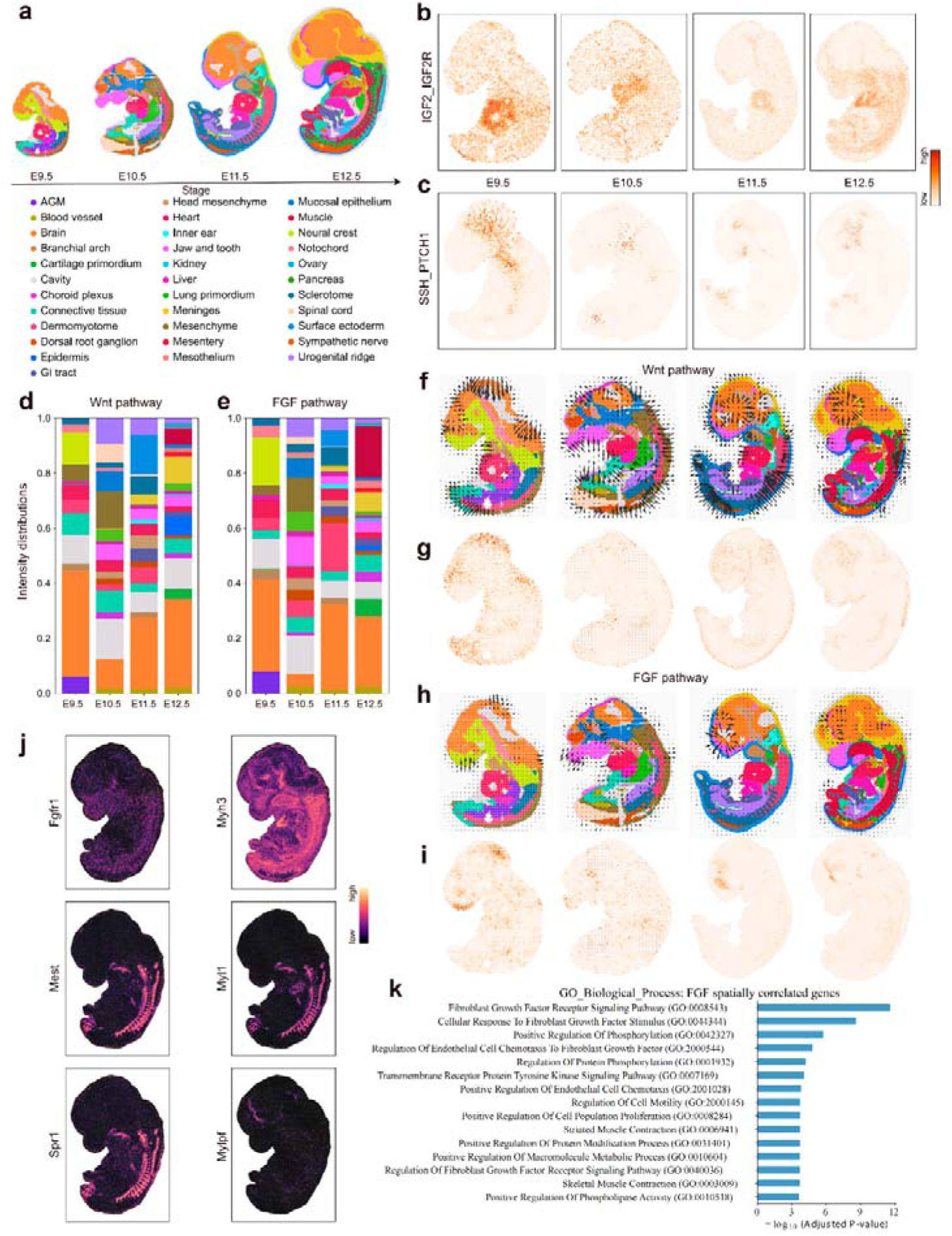
| SpaGRD reveals spatial-temporal ligand–receptor interactions and pathway activities during mouse embryonic development. **a**, Region annotation information of mouse embryo sections from stage E9.5 to E12.5, annotated by tissue types based on the original study. **b**, Spatial intensities of IGF2–IGF2R interaction inferred by SpaGRD, exhibiting high intensity in multiple regions, particularly in the heart. **c**, Spatial intensities of SHH–PTCH1 interaction, exhibiting localized intensity in brain and gastrointestinal (GI) regions. **d**, Regional intensity distribution of Wnt pathway activities across four developmental stages. **e**, Regional intensity distribution of FGF pathway activities across four developmental stages. **f**, Vector fields indicating the spatial flow of inferred Wnt signaling pathway. **g**, Spatial intensities of the Wnt signaling pathway, highlighting its activity along the dorsal side and in the brain. **h**, Vector fields indicating the spatial flow of inferred FGF signaling pathway. **i**, Spatial intensities of the FGF signaling pathway, showing strong activity in the brain and muscle regions across developmental stages. **j**, Expression patterns of selected genes that are either components or targets of the FGF signaling pathway, such as *Fgfr1*, *Mest*, *Spp1*, *Myh3*, *Myh1*, and *Mylpf*, revealing both global and tissue-specific correlations with FGF signaling. **k**, GO enrichment analysis of genes positively correlated with FGF signaling activity, showing terms related to FGF receptor signaling, cell proliferation, and muscle development.

Next, we investigated the characters of pathway-level interactions. Our results revealed that the Wnt signaling pathway, which plays a crucial role in regulating body patterning and organogenesis, exhibited strong activity across multiple regions and developmental stages (**Fig. 6d**). Notably, elevated Wnt signaling was observed in the brain and dorsal regions, including the surface ectoderm at E11.5 and the epidermis at E12.5 (**Fig. 6g**), consistent with previous findings^43–46^. Spatially, Wnt pathway interactions along the dorsal axis extended from the external surface inward, suggesting a role in orchestrating the development of deeper tissue layers (**Fig. 6f**). The FGF signaling pathway, known for its regulation of cell proliferation, migration, and differentiation during embryogenesis^47–49^, also showed widespread activity. Strong FGF signaling interactions were observed in various organs, particularly in the brain and muscle, across multiple stages of development (**Fig. 6h** and **Fig. 6i**). Interestingly, although both Wnt and FGF pathways are active in the brain, they exhibit distinct spatial patterns, indicating that each may contribute to the formation of different brain structures. In addition to Wnt and FGF, we also analyzed TGFβ and SEMA3 signaling pathways. Both exhibited unique spatial interaction patterns that are likely integral to specific aspects of embryonic development (**Supplementary Fig. 11a–f**), further emphasizing the pathway-specific spatial regulation of developmental processes.

Finally, we investigated the functional relationships between four signaling pathways and their associated genes. Using FGF signaling as a representative example, we performed correlation analysis between FGF signaling intensity distribution and gene expression profiles, employing the Pearson correlation coefficient to identify positively correlated genes. Among these, some genes were ligands or receptors of the FGF pathway, such as *Fgfr1* (**Fig. 6j**), while others were known FGF-responsive genes, such as *Mest*^50–52^. Notably, the expression pattern of *Mest* closely matched the FGF signaling intensity (Fig. 6j). However, several positively correlated genes showed only localized concordance between their expression and pathway activity rather than a global correspondence. For example, *Spp1* exhibited higher expression in brain regions, whereas *Myh3*, *Myh1*, and *Mylpf* were predominantly expressed in muscle tissue (**Fig. 6j**), suggesting these genes respond to FGF signaling in a tissue-specific or context-dependent manner. This observation likely reflects the functional diversity of FGF signaling during development. Additionally, GO analysis of positively correlated genes for each signaling pathway across four developmental stages revealed enrichment patterns consistent with the known biological roles of these pathways (**Fig. 6k** and **Supplementary Fig. 12a–c**).

## Discussion

Cell-cell communication is a critical application in spatial transcriptomics, attracting significant attention due to its importance in understanding fundamental biological processes. Although numerous methods have been developed, particularly those tailored to spatial transcriptomic data, which emphasize the fact that intercellular interactions typically occur within limited spatial distance, most current approaches rely heavily on manually defined rules or attention-based deep learning architectures. However, such strategies often introduce excessive bias, as they are not grounded in first principles. To address this limitation, there is an urgent need for a biophysically meaningful model that captures the extracellular diffusion and reaction dynamics of ligand–receptor pairs. While several theoretical studies have attempted to construct such models to reveal the complexity of these dynamics, they often involve overly stringent assumptions and highly intricate formulations, making them impractical for direct application to real-world spatial transcriptomics data. As a result, these models largely remain theoretical in nature. We argue that an effective mathematical framework for inferring cell-cell communication from spatial omics data should satisfy two key criteria: (1) It must be grounded in realistic biophysical principles, accurately reflecting the nature of ligand–receptor interactions; (2) It must be practically solvable and applicable to spatial transcriptomics datasets, which may require appropriate simplifications to balance realism and tractability.

In this study, we propose SpaGRD, a first-principles-based framework that models cell-cell communication by simulating the extracellular diffusion of ligands and their reactions with receptors using Fick’s law and the law of mass action. This formulation establishes a biophysical dynamical system that captures the essential processes of ligand transport and binding. To accommodate various spatial transcriptomics technologies, SpaGRD leverages GSP to represent cellular entities, spatial coordinates, and gene expression profiles on irregular spatial domains. By efficiently solving the proposed dynamical system, SpaGRD infers the spatial distribution of ligand–receptor reaction intensities. Moreover, downstream analyses based on GSP enable the identification of ligand–receptor interaction modules with heterogeneous spatial patterns, achieving molecular-level insights at single-cell resolution. Additionally, by adopting a divide-and-conquer strategy, SpaGRD traces the reaction-diffusion trajectory of ligands secreted by individual cells, allowing the reconstruction of a cell-cell communication network. This network can be visualized as a vector field, illustrating the directionality and strength of intercellular communication, thereby enabling cell-level inference at single-cell resolution.

We first validated the performance of SpaGRD on simulated datasets. Notably, the graph-based reaction-diffusion equations can be regarded as a numerical solver for PDEs, similar to the finite difference method. Both approaches yield highly comparable results. However, the graph-based formulation offers greater flexibility, allowing for arbitrary spatial distributions of spots or cells. More importantly, graphs enable the representation of spots or cells as nodes, thereby facilitating the modeling of signal flow and interactions between spatial units. Further, we applied SpaGRD to multiple real datasets. In mouse brain data, we revealed ligand–receptor dynamics and identified spatially coherent ligand–receptor modules. In brain metastasis datasets, we uncovered spatially distinct responses of COL1A1–CD44 and COL6A1–CD44, demonstrating that ligand–receptor interactions can provide insights into intratumoral heterogeneity. In human colorectal cancer data, SpaGRD characterized the spatial molecular signatures of key pathways within tumor tissues. In mouse embryonic data, we captured spatial distribution patterns and dynamic changes in cell-cell communication during development. Together, these applications demonstrate the broad utility of SpaGRD in dissecting cell-cell communication across diverse spatial omics contexts.

We deliberately simplified the ligand–receptor interaction process to ensure computational tractability. However, real biochemical signaling is considerably more complex^53^. Factors such as dynamic changes in cell positioning, dissociation of ligand–receptor complexes, and the influence of cell morphology on diffusion are not yet captured in our current model. Future work should aim to incorporate these biological details to develop more refined and accurate representations of cell-cell communication. Moreover, SpaGRD focuses solely on the ligand–receptor binding process. In reality, this interaction is embedded within a broader signaling cascade, involving upstream ligand production and downstream intracellular signaling events. Future models could benefit from integrating these additional layers to better represent the full communication process. Ultimately, by leveraging first-principles-based modeling, we believe that cell-cell communication inference in spatial omics will become more mechanistic, interpretable, and biologically grounded.

## Methods

### Data Preparation and Initialization

#### Spatial transcriptomics preprocessing

While intercellular ligand–receptor interactions are fundamentally mediated by protein-level dynamics, measuring transcriptome-wide spatial protein concentrations remains technically challenging. Therefore, consistent with established practices in the field, we utilize spatially resolved RNA expression as a practical and biologically informative proxy for local protein abundance. Raw count matrices were first filtered to remove lowly expressed genes and normalized to counts per million (CPM), followed by log-transformation. Crucially, to satisfy the mathematical constraints of the law of mass action utilized in our kinetic models, we applied an additional global scaling step. The normalized count matrix was scaled such that its maximum value equals one, ensuring that these simulated molecular concentrations—acting as computational surrogates for actual protein levels—remain physically interpretable and computationally stable.

#### Ligand–receptor interaction database

We mapped spatial gene expression to potential communication events using the CellChatDB database, which provides curated ligand–receptor interactions and extensive metadata. Furthermore, to accommodate specialized biological contexts and novel discoveries, SpaGRD is highly extensible and fully supports custom, user-defined ligand–receptor pairs. Because reaction-diffusion dynamics inherently model the movement of molecules through extracellular space, we strictly filtered the database to exclude contact-dependent interactions. Our analysis focuses exclusively on secreted signaling and extracellular matrix (ECM) interactions, as spatial transcriptomic measurements of these genes translate to diffusible proteins that fit the continuous spatial diffusion assumption. For multimeric ligands or receptors, SpaGRD conservatively estimates the functional complex abundance by taking the minimum expression level among its constituent subunits.

### Physical intuition and continuous PDEs

Cell-cell communication via secreted factors is driven by two coupled physical processes: the diffusion of ligands across the tissue architecture and their localized binding to cognate receptors. Let *u_i_*(***x***, *t*) and *v_j_*(***x***, *t*) denote the concentration of ligand *i* and receptor *j* at spatial coordinate 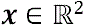 (or 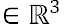) and time 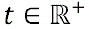 According to Fick’s law, ligands move down their concentration gradient, described by the Laplacian operator Δ*u_i_*(*x*, *t*) mathematically. Simultaneously, according to the law of mass action, the binding event occurs proportionally to the product of the available ligand and receptor concentrations, *u_i_*(***x***, *t*) × *v_j_*(***x***, *t*).

Because a single ligand can often bind multiple receptors and vice versa, SpaGRD establishes a generalized dynamic system to model these complex many-to-many competitive relationships. Let *P_i_* denote the set of receptors interacting with ligand *i*, and *Q_i_* denote the set of ligands interacting with receptor *j*. The continuous reaction-diffusion process is governed by the following system of partial differential equations (PDEs):

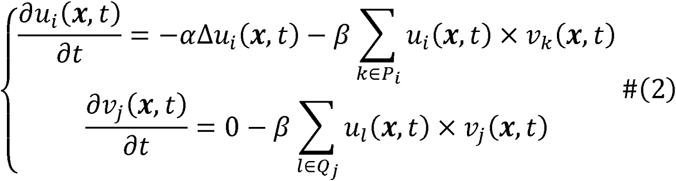

Here, α parameterizes the diffusion rate, controlling how far signals travel, while β represents the reaction rate governing binding affinity.

### Transformation to graph signals

Solving continuous PDEs analytically is intractable for spatial omics data because biological tissues are captured as discrete, non-uniform spatial spots rather than regular continuous grids. To bridge this gap, SpaGRD transforms the physical equations into the domain of Graph Signal Processing (GSP).

Denote *n* as the number of cells/spots in spatial transcriptomics, and the spatial graph can be constructed and represented as *G* = (*V*, *E*) where *V* and *E* are the node set (related to cells/spots) and edge set, respectively. An edge between two cells/spots can be defined by K-Nearest Neighbors (KNN) or Radius Nearest Neighbors (RNN). And the adjacency matrix 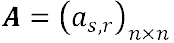 of the graph is defined as:

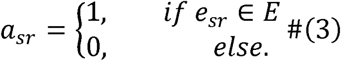

where *e_sr_* is the edge between spot/cell *s* and spot/cell *r* and *n* is the number of spots/cells. The degree matrix *D* = *diag*(*d*_1_, *d*_2_, … *d_n_*) of graph *G*, in which 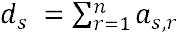, represents the degree of the *s*-th node calculated by the sum of the number of edges connecting at the node. In this way, the Laplacian matrix is defined as:

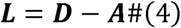

Once the spatial graph is constructed, the ligand/receptor concentration functions can be transformed from continuous form to discreate form:

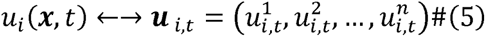

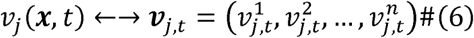

where 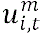 and 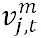 are the ligand concentration and receptor concentration on cell/spot *m* and time *t*, respectively and m = 1,2, …, *n*. The double-arrow notation indicates that the left and right formulas represent the same quantity of molecules under two equivalent formulations: the continuous mathematical representation (left) and the corresponding graph-signal representation on the constructed graph (right). In this way, ***u****_i,t_* and ***v****_j,t_* can be treated as the graph signals on the graph. And the corresponding discrete forms of diffusion and reaction are:

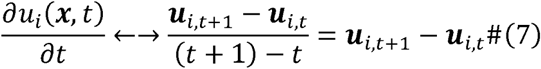

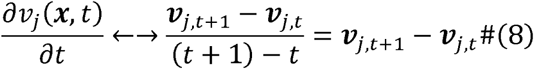

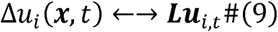

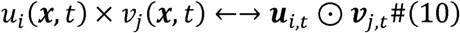

where ⊙ is the Hadamard product. So far, we have obtained discrete forms of terms in diffusion-reaction equations on the graph.

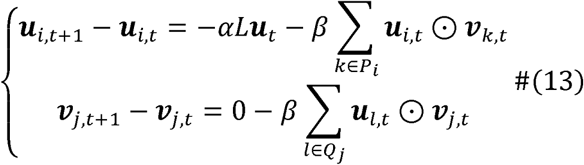

After the proper adjustment, one has the molecule changes of the dynamical system along time *t* to *t*+1:

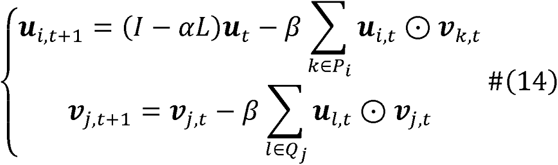

### Molecular-centric analysis

#### Ligand-receptor interaction inference

Based on above transformation, the most important item, ligand–receptor interactions 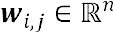 can be obtained easily by summing the interactions of all time steps:

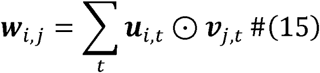

where ***w****_i,j_* is an *n*-dimensional vector, showing the ligand–receptor interaction intensity distributions on each cell/spot for a given ligand–receptor pair, and *n* is the number of spots. A biological pathway may contain multiple ligand–receptor pairs. Hence, we can merge the interactions from individual ligand–receptor interaction pairs to produce pathway-level interaction inference 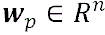:

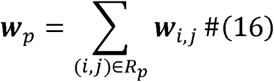

where *R_p_* is the set of ligand–receptor interaction pairs that belong to the same pathway *p*.

#### Spatially variable interaction selection and clustering

The ligand–receptor interaction vector can be viewed as a graph signal on spatial graph. Hence, we implement graph signal processing analysis based on graph Fourier transform similar to our previous work, SpaGFT. Briefly, ***w****_i,j_* is the representation, which neglects the spatial topology information among the cells/spots. By implementing graph Fourier transform, we obtain the novel frequency-domain representation, where each component of the frequency-domain signal corresponds to a different frequency. And low-frequency is related to smooth and well-structured patterns while high-frequency is related to noise or severe fluctuations on graph. Hence, we stress the importance of low-frequency components by giving high weights to determine which ligand–receptor interaction pairs exhibit a spatial specificity pattern. In addition, ligand–receptor pairs with similar spatial patterns were identified by unsupervised clustering algorithm based on frequency-domain representations. Here, we adopt Louvain algorithm to obtain ligand–receptor (LR) modules, defined as ligand–receptor pair sets with consistent spatial patterns. Moreover, spots with high interaction activities corresponding to a LR module were picked out based on *k*-means algorithm with *k* = 2, to represent the LR module’s region. More details about the frequency-domain analysis can be found in Supplementary Note.

#### Gene-interaction correlation analysis

To investigate the relationships between genes and interactions, we performed correlation analysis between interaction vectors and gene expression profiles. For each pathway or ligand–receptor (LR) module, an interaction vector was computed by aggregating the interactions of all constituent ligand–receptor pairs. We then calculated the Pearson correlation coefficient between this interaction vector and the expression of each gene. Genes showing high correlation with the interactions (default cutoff of 0.2) were selected for downstream functional analyses, such as GO enrichment analysis.

#### Growth rate analysis

SpaGRD can infer the overall ligand–receptor interaction intensity at each time point *t*’:

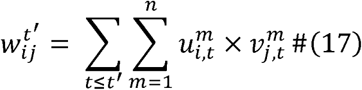

We also utilize this to determine whether the system has reached a steady state when the 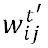 achieves a stable development trend at time *t*’. SpaGRD provides an incremental curve plot for users to determine the number of steps in a dynamical system. By combining interaction intensity charts across different time points, we illustrate dynamical growth trends over time.

### Cell-centric analysis

#### Cell-cell interaction scoring

While the above analysis shows the interaction intensity on all cells/spots, we are also interested in which cells interact with surrounding cells. To achieve this, we adopted a divide-and-conquer strategy due to the property of linear additivity. For ligand 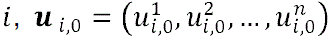 is the concentration distribution on *n* cells/spots initially, described by normalized gene expressions. SpaGRD splits *u*_*i*,0_ into a set of vectors 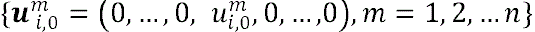. Each 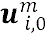 is related to initial ligands on cell/spot *m*. These 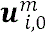 will interact with receptors as independently initial graph signals to acquire the status 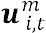 at time *t* and interactions. And solve the dynamic system to produce:

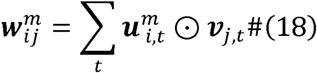

where 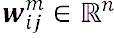. It is obvious that interactions 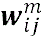 must reflect the interactions of cell/spot *m* to others because only cell/spot *m* contains ligand molecules. Hence, we can obtain cell-cell interaction in this way:

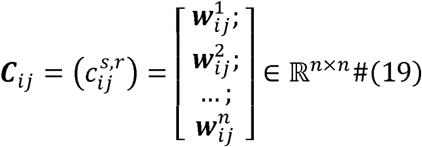

where 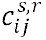 indicates the interactions from cell/spot *s* to cell/spot *r* under ligand *i* and receptor *j*.

#### Cluster-level analysis and interaction direction analysis

If spot/cell labels are known beforehand, interactions from cells or spots sharing the same label can be averaged or summarized to obtain cell-type–level interaction information. To identify statistically significant interacting cell types, a permutation test is performed by randomly rearranging labels 1,000 times (default), with significance determined at *p*-value□<□0.05. SpaGRD also offers visualization options, including chord diagrams and network plots. To visualize signaling directions, we implement field vector analysis based on *C_i,j_* for sender or receiver cells. Briefly, the interaction vector at each location is converted into a two-dimensional vector to indicate the direction of signaling, following the approach described in the COMMOT paper.

### Downstream analysis based on graph ligand-receptor interaction system

#### Detection of spatially variable ligand–receptor interactions

To identify ligand–receptor interaction pairs with distinct spatial patterns in SpaGRD, we integrated our previously developed tool, SpaGFT, which leverages graph signal processing to detect spatially variable molecular signals. Briefly, SpaGFT constructs a spatial graph and performs spectral decomposition to derive Fourier modes spanning low to high frequencies. The interaction vector of each ligand–receptor pair is treated as a graph signal and transformed into the frequency domain to obtain its Fourier coefficients. These coefficients are then weighted according to their corresponding frequencies to calculate a *GFTscore*, which quantifies the spatial variability of the interaction. Ligand–receptor pairs exhibiting pronounced spatial patterns are subsequently identified based on their ranked *GFTscore*s.

#### Ligand–receptor module (LR module) identification

Some ligand–receptor interaction pairs may exhibit similar spatial patterns, suggesting coordinated signaling events. To identify such coordinated interactions, we perform interaction module analysis via clustering. Unlike conventional approaches that rely solely on individual interaction profiles, our method integrates both spatial context and interaction intensity by leveraging the representational power of SpaGFT. Specifically, we compute frequency-domain representations of ligand–receptor pairs by combining interaction intensity with the underlying spatial graph. Clustering is then conducted on the low-frequency components of these representations using the Leiden algorithm, grouping ligand–receptor pairs into spatial interaction modules. To visualize the clustering results, we apply UMAP to the frequency-domain features of the ligand–receptor pairs. For each identified module, we further determine the cells or spots exhibiting high interaction activity related to the module’s ligand–receptor pairs. This is achieved by applying *k*-means clustering (*k* = 2) to distinguish between spots with high versus low interaction levels.

#### Correlation analysis between LR modules and gene expression

For each ligand–receptor interaction module, we first aggregate the interaction signals of all ligand–receptor pairs within the module to generate a representative module-level interaction profile. We then compute the Pearson correlation coefficient (PCC) between this profile and the expression level of each gene across spatial locations. Genes with a positive correlation (PCC > 0.2) are positively associated with the ligand–receptor interaction module.

#### Correlation analysis between pathway signaling and genes

For a given signaling pathway, we aggregate the interaction signals of all associated ligand–receptor pairs to construct a representative pathway-level interaction profile. We then compute the Pearson correlation coefficient (PCC) between this profile and the expression level of each gene across spatial locations. Genes with a positive correlation (PCC > 0.2) are positively associated with the signaling activity of that pathway.

### Benchmarking settings

#### Simulation dataset generation

Due to the complexity of ligand–receptor interactions, where a single ligand can bind to multiple receptors and vice versa, we designed ten simulation scenarios to capture a diverse range of interaction patterns, including one-to-one, one-to-many, and many-to-many relationships. For each scenario, 10 replicates were generated. The expression profiles of ligands and receptors were simulated using simstpy^19^, a spatial covariance-based computational tool previously developed for simulating spatial transcriptomics data. To obtain ground truth distributions of interaction intensities across spatial locations, we applied the standard finite difference method (FDM), a widely used numerical approach for solving PDEs. This enabled us to compute the interaction strength between ligand–receptor pairs at each spatial spot under the specified interaction models. Additionally, one extra replicate was generated for each simulation case to serve as a validation set for parameter tuning across all evaluated methods.

#### Metrics used for evaluation

To measure the accuracy in predicting ligand–receptor interaction, two metrics, PCC and RMSE, were included, where PCC measures the linear correlation and RMSE measures the estimation error between the estimated interaction values and the ground truth. For a ligand–receptor pair, PCC is defined as

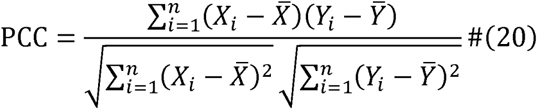

and MSE is defined as

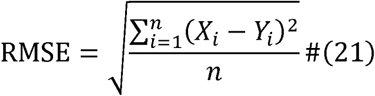

where *Y* is ground truth values, *X* represents the predicted ligand–receptor interactions. To overcome the scale effect, the predicted ligand–receptor interactions were normalized to enable the same maximum values as the ground truth for each competing method.

### Parameter setting of other tools

1. COMMOT (version 0.0.3). COMMOT is an optimal transport model that simultaneously considers the competitive relationships between multiple ligand-receptor pairs and the spatial distance between cells. We follow the tutorial and run *ct.tl.spatial_communication* to obtain the diffusion matrix, which is stored in *adata-dis500.obsp*. To obtain results consistent with SpaGRD, we first sum the diffusion matrix of the ligand by row and then multiply it with the receptor to obtain the reaction intensity. We adjusted three parameters, *cot_ eps_p* [0.05, 0.1, 0.2], *dis_thr* [30, 50, 100], and *cot_rho*[5, 10, 15], generating 27 parameter combinations, on parameter tuning datasets. And the parameter combination with best performance were used on independent datasets in benchmarking results.
2. Spacia. Spacia models the expression of the one receiving cell (as the response variable) as the function of the expression signature of multiple sending cells in the vicinity of this receiving cell, using a MIL approach. We first follow its tutorial to obtain *Interactions.csv*. In order to unify the results with SpaGRD, we use the *Primary_instance score* of the signals received by each spot to calculate the interaction strength with the Receiver. We adjusted several parameters, *receiver_features*/*sender_features* [5, 10, 15], *response_exp_cutoff* [’auto’, 0.33, 0.67], and *mcmc_params*[(50000, 25000, 10, 3), (25000, 12500, 5, 2), (12500, 6250, 3, 1)], generating 27 parameter combinations, on parameter tuning datasets. And the parameter combination with best performance were used on independent datasets in benchmarking results.
3. SpatialDM (version 0.2.0). SpatialDM is a statistical model which uses a bivariant Moran’s statistic to detect spatially co-expressed ligand and receptor pairs, their local interacting spots, and communication patterns. According to its tutorial, we firstly construct weight matrix using *sdm.weight_matrix* function and then run *sdm.spatial_local* and *sdm.sig_spots* to calculate cell-cell communication patterns. Note that SpatialDM uses *p*-value to represent interaction intensity. Hence, we use one minus the *p*-value to obtain the SpatialDM’s prediction results. In addition, we used *sdm.spatial_local* because we also tried *sdm.spatial_global*, which exhibited lower performance. One parameter was picked out for parameter selection due to limited adjustment space: *l* [1 to 27] in *sdm.weight_matrix*, generating 27 parameter settings, on parameter tuning datasets. And the parameter combination with best performance was used on independent datasets in benchmarking results.
4. SpaGRD. SpaGRD infers cell-cell communication based on the constrction of graph reaction-diffusion equations and solves the equations using graph signal processing techniques. We run *SpaGRD.grd.reaction_diffusion_system* to obtain the ligand–receptor interaction results directly after processing. Three parameters were picked out for parameter selection: *alpha* [0.25, 0.5, 0.75], *beta* [0.25, 0.5, 0.75], *step* [2000, 3000, 4000] in *SpaGRD.grd.reaction_diffusion_system* function, generating 27 parameter combinations, on parameter tuning datasets. And the parameter combination with best performance was used on independent datasets in benchmarking results.

### Case study implementation details

#### Identification of spatial domain and cell type in human colorectal cancer data

We applied the Leiden algorithm to identify spatial domains in the Visium data. Given the scale of VisiumHD datasets, rapid annotation was performed by calculating the Pearson correlation coefficient (PCC) between spots of VisiumHD data and cells from matched scRNA-seq references data with annotation.

#### Analysis of cell type colocalization in human colorectal cancer data

For each cell, we computed the proportion of surrounding cell types among its eight nearest spatial neighbors. The average of these neighboring proportions was calculated for cells sharing the same cell type, generating a robust metric for spatial colocalization intensity.

## Code availability

The SpaGRD software is developed in Python. All custom code, scripts, and tutorials required to reproduce the analyses presented in this study will be deposited and made publicly accessible on GitHub under the MIT license immediately upon publication.

## Data availability

The publicly available datasets used in this study can be accessed from the following sources: mouse brain 10x Visium dataset (https://www.10xgenomics.com/datasets/mouse-brain-section-coronal-1-stand ard); Xenium brain metastasis dataset (https://zenodo.org/records/10712720); human colorectal cancer VisiumHD datasets (https://www.10xgenomics.com/products/visium-hd-spatial-gene-expression/d ataset-human-crc); human colorectal cancer Visium dataset (https://www.10xgenomics.com/datasets/human-colorectal-cancer-whole-tran scriptome-analysis-1-standard-1-2-0); mouse embryo Stereo-seq datasets (https://db.cngb.org/stomics/mosta/).

## Acknowledgement

This work was supported by National Key R&D Program of China [2020YFA0712400]; National Nature Science Foundation of China [NSFC, 62272270]; Open-project of BGI-ShenZhen, Shenzhen 518000, China [BGIRSZ20220005]; Shandong Provincial Natural Science Foundation for Distinguished Young Scholars [ZR2023JQ002]. We also thank Dr. Qin Ma and Dr. Yuzhou Chang from Ohio State University for their valuable support and discussions on methodology.

## Author contributions

B.L. conceptualized and supervised the project. J.L. and S.S. designed and implemented the method. S.S. and J.L. implemented simulation and benchmarking. S.S. and J.L. collected the data. J.L., S.S., Z.C., Z.L, B.L. and G.L. interpreted the results in case studies and wrote the paper. All authors reviewed the manuscript.

## Supplementary Figures

**Supplementary Fig. 1.**
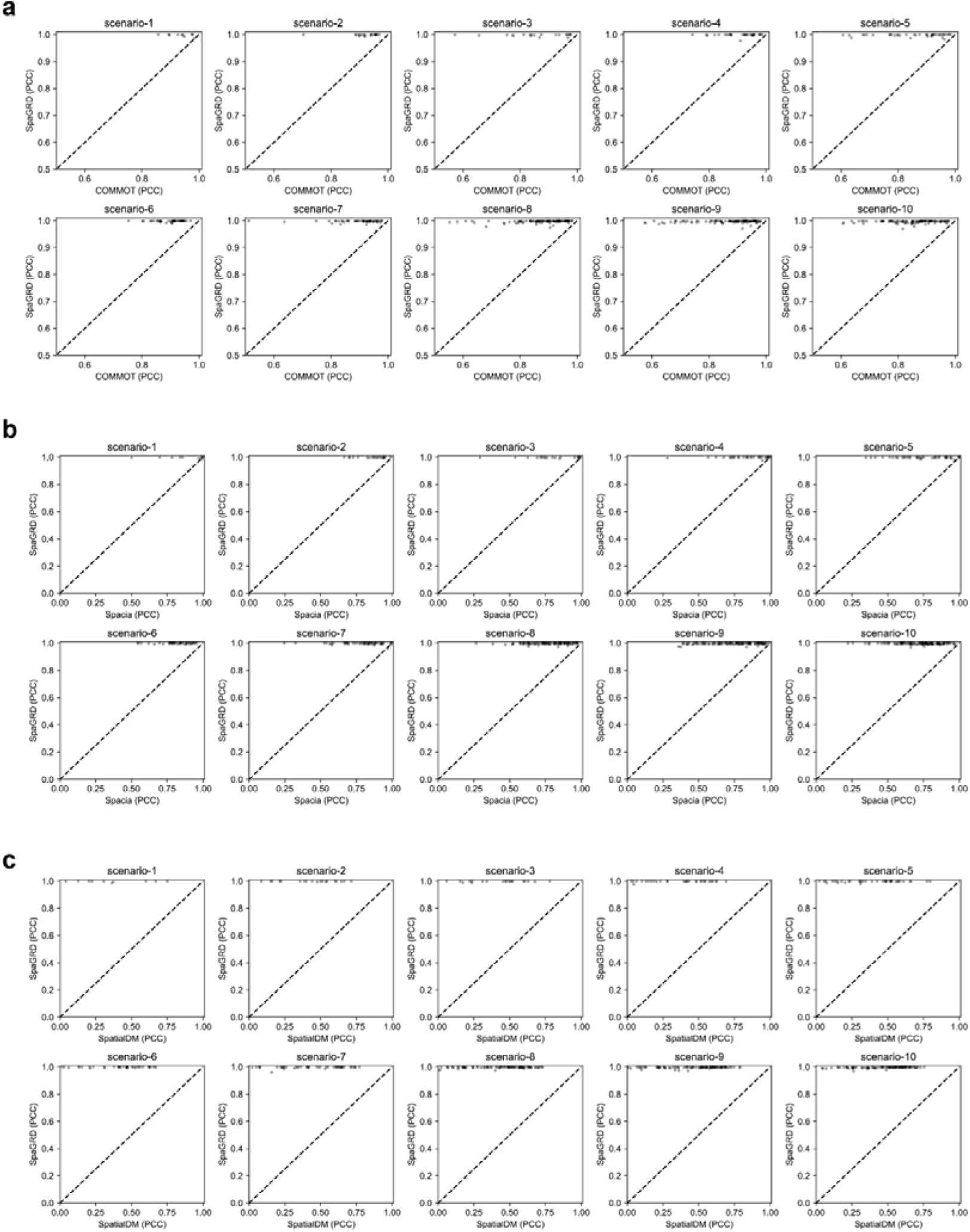
| The pair wise performance comparisons using PCC. **a**, The performance comparisons between SpaGRD and COMMOT on 10 simulation scenarios under PCC. Each point represents a ligand-receptor pair in a replicate. **b**, The performance comparisons between SpaGRD and Spacia on 10 simulation scenarios under PCC. Each point represents a ligand-receptor pair in a replicate. **c**, The performance comparisons between SpaGRD and SpatialDM on 10 simulation scenarios under PCC. Each point represents a ligand-receptor pair in a replicate.

**Supplementary Fig. 2.**
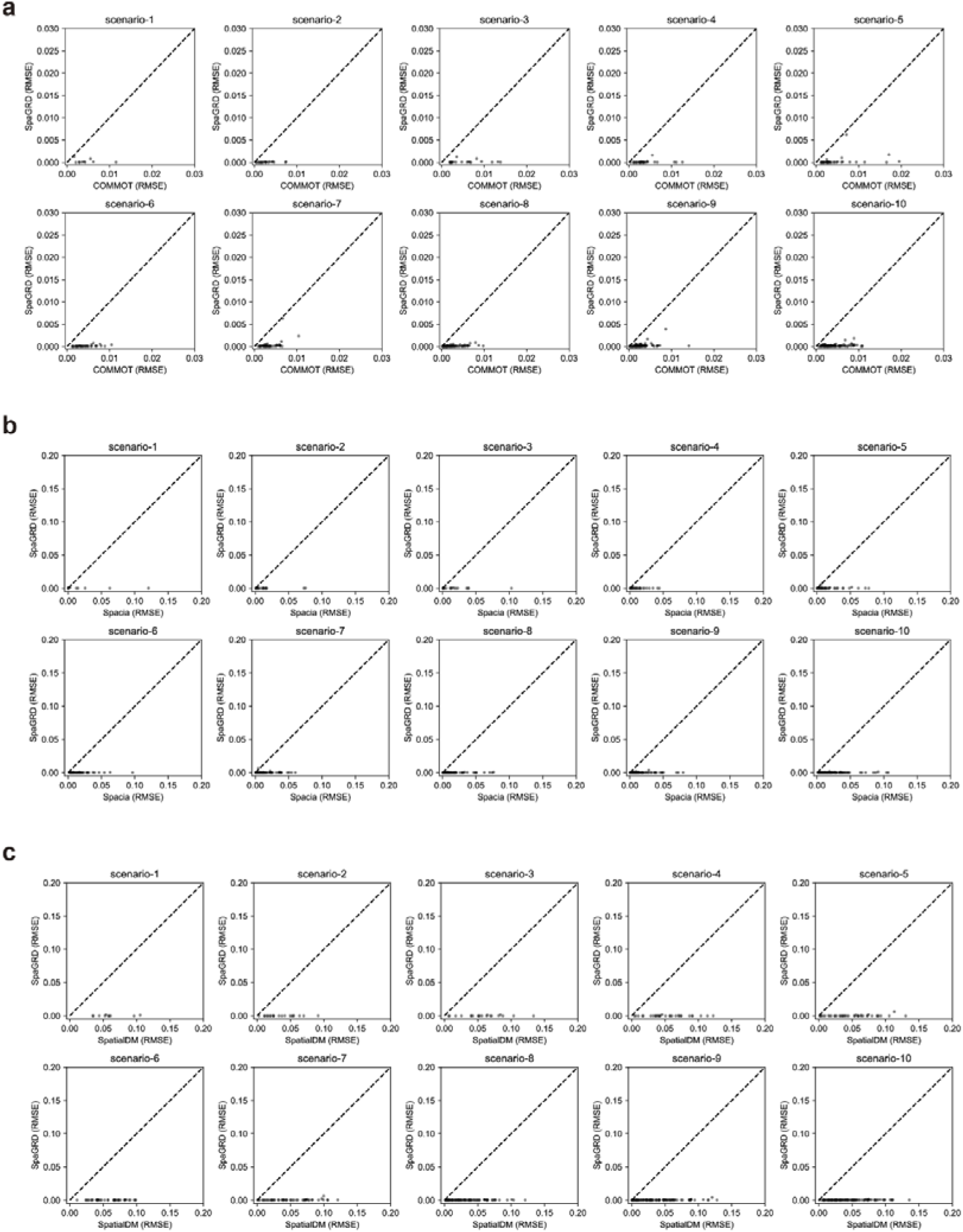
| The pair wise performance comparisons using RMSE. **a**, The performance comparisons between SpaGRD and COMMOT on 10 simulation scenarios under RMSE. Each point represents a ligand-receptor pair in a replicate. **b**, The performance comparisons between SpaGRD and Spacia on 10 simulation scenarios under RMSE. Each point represents a ligand-receptor pair in a replicate. **c**, The performance comparisons between SpaGRD and SpatialDM on 10 simulation scenarios under RMSE. Each point represents a ligand-receptor pair in a simulation replicate.

**Supplementary Fig. 3.**
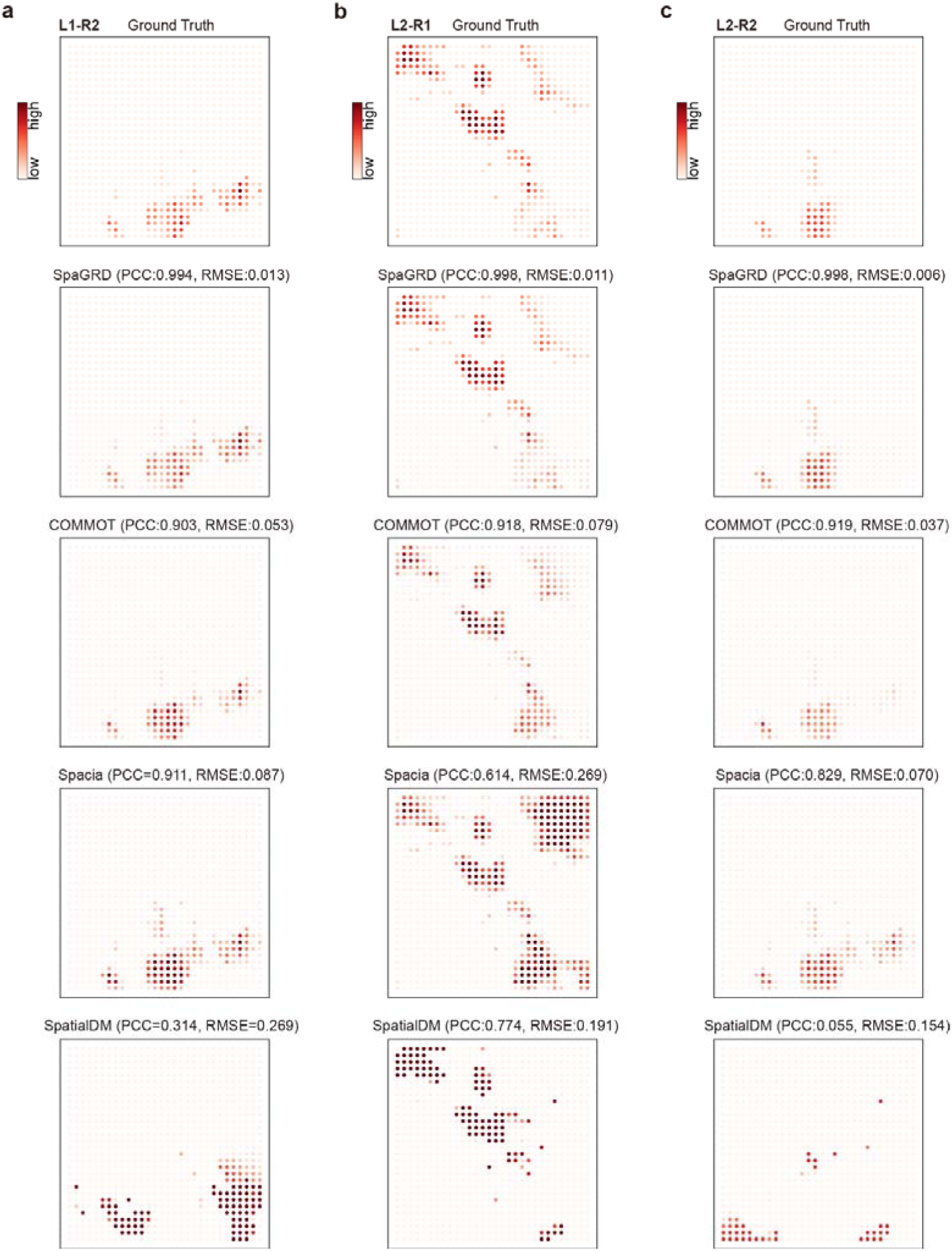
| Examples of ligand-receptor interaction predictions. Ground truth interaction intensity between ligand and receptor, obtained by numerical simulation of PDEs. Predicted interaction intensities between ligand and receptor by four methods: SpaGRD, COMMOT, Spacia, and SpatialDM. Each panel includes the Pearson correlation coefficient (PCC) and root mean square error (RMSE) with respect to the ground truth. Color intensity indicates the predicted strength of interaction at each spatial location. **a**, The comparisons on ligand L1 and receptor R2. **b**, The comparisons on ligand L2 and receptor R1. **c**, The comparisons on ligand L2 and receptor R2.

**Supplementary Fig. 4.**
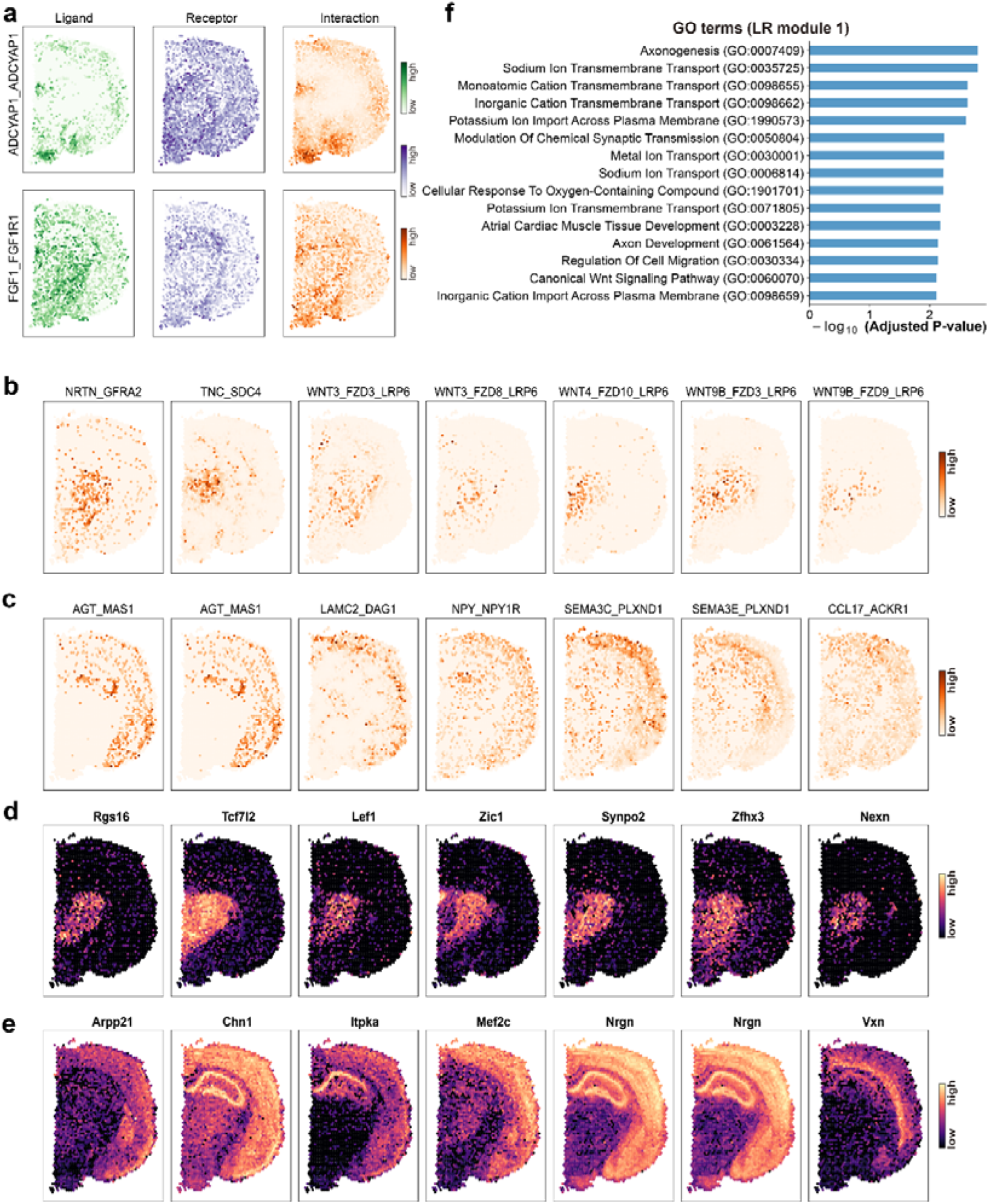
| Spatial interaction modules and their associated gene expression and functional biological terms. **a**, Spatial expression patterns of ligands, receptors, and inferred interaction intensities for two representative ligand–receptor pairs: ADCYAP1–ADCYAP1R1 and FGF1–FGFR1, showing distinct spatial overlap and interaction regions. **b**, Spatial distributions of representative ligand–receptor interaction pairs in LR module 1, including NRTN_GFRA2, TNC_SDC4, WNT3_FZD3_LRP6, WNT3_FZD8_LRP6, WNT4_FZD10_LRP6, WNT9B_FZD3_LRP6, and WNT9B_FZD9_LRP6. These interactions are primarily enriched in the thalamus region. **c**, Representative ligand–receptor interaction pairs from LR module 2, including AGT_MAS1, LAMC2_DAG1, NPY_NPY1R, SEMA3C_PLXND1, SEMA3E_PLXND1, and CCL17_ACKR1, which are mostly localized in the cortex region. **d**, Spatial expression of genes positively correlated with LR module 1, such as Rgs16, Tcf7l2, Lef1, Zic1, Synpo2, Zfhx3, and Nexn, showing similar spatial enrichment to the interaction pairs in module 1. **e**, Spatial expression of genes correlated with LR module 2, including Arpp21, Chn1, Itpka, Mef2c, Nrgn, and Vxn, showing distinct spatial localization in the cortex. **f**, Gene Ontology (GO) enrichment analysis of genes positively correlated with LR module 1. Enriched terms include axonogenesis, canonical Wnt signaling pathway, and various ion transmembrane transport processes, suggesting that the interaction module is involved in neurodevelopmental and electrophysiological processes.

**Supplementary Fig. 5.**
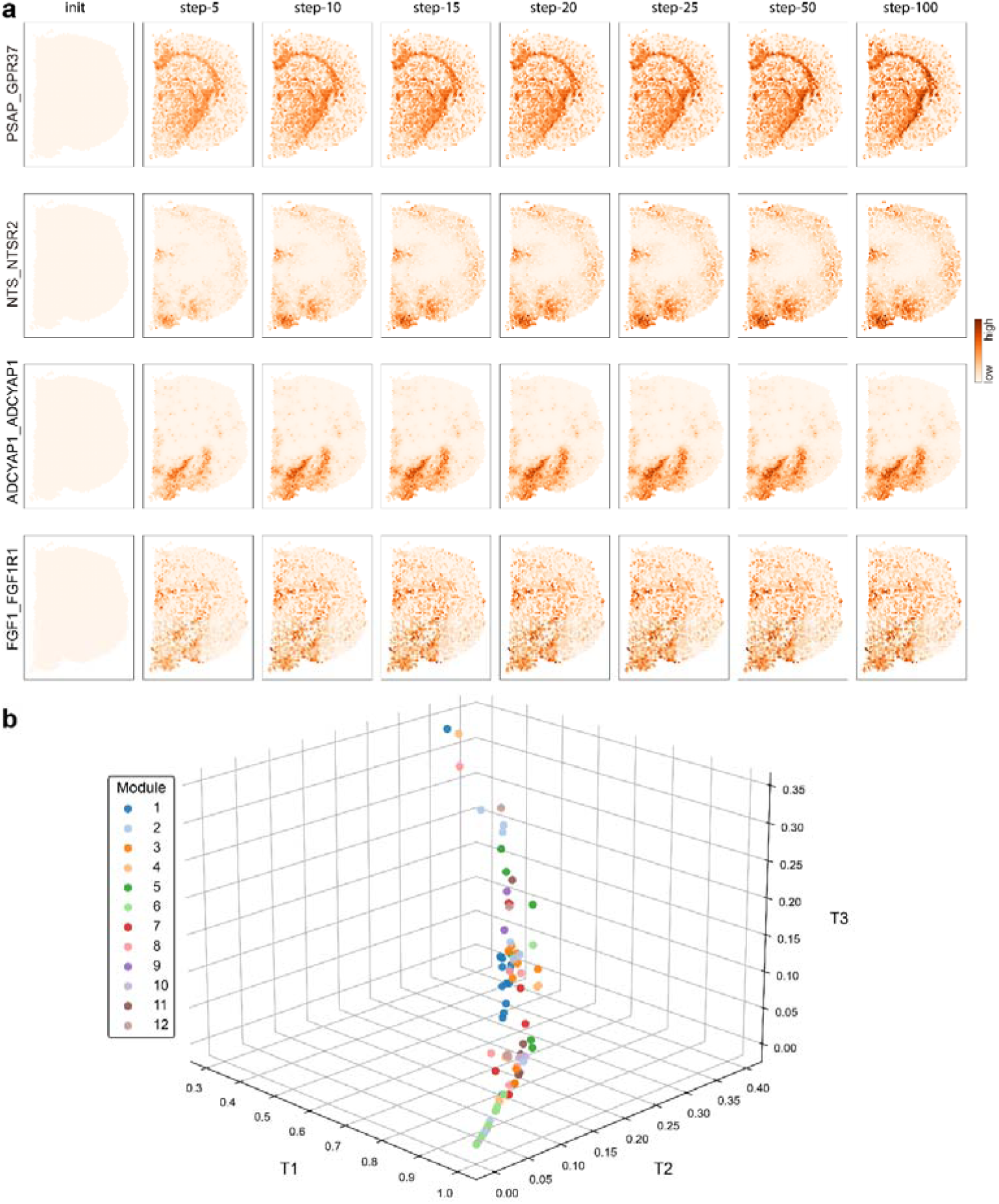
| Dynamic modeling of ligand–receptor interaction intensity reveals temporal diversity in communication patterns. **a**, Time-resolved interaction maps for four ligand–receptor pairs (PSAP–GPR37, NTS–NTSR2, ADCYAP1–ADCYAP1R1, and FGF1–FGFR1) across ten simulation steps, from initialization to step 100. The results show different temporal dynamics, with some interactions forming rapidly and others gradually intensifying over time. **b**, Three-dimensional scatter plot representing the proportion of total interaction intensity occurring in the three step intervals: T1 (steps 0–5), T2 (steps 6–30), and T3 (steps 31–100). Each dot represents a ligand–receptor pair, colored by module identity. The distribution reveals variability in interaction speed and dynamics across different pairs and signaling modules.

**Supplementary Fig. 6.**
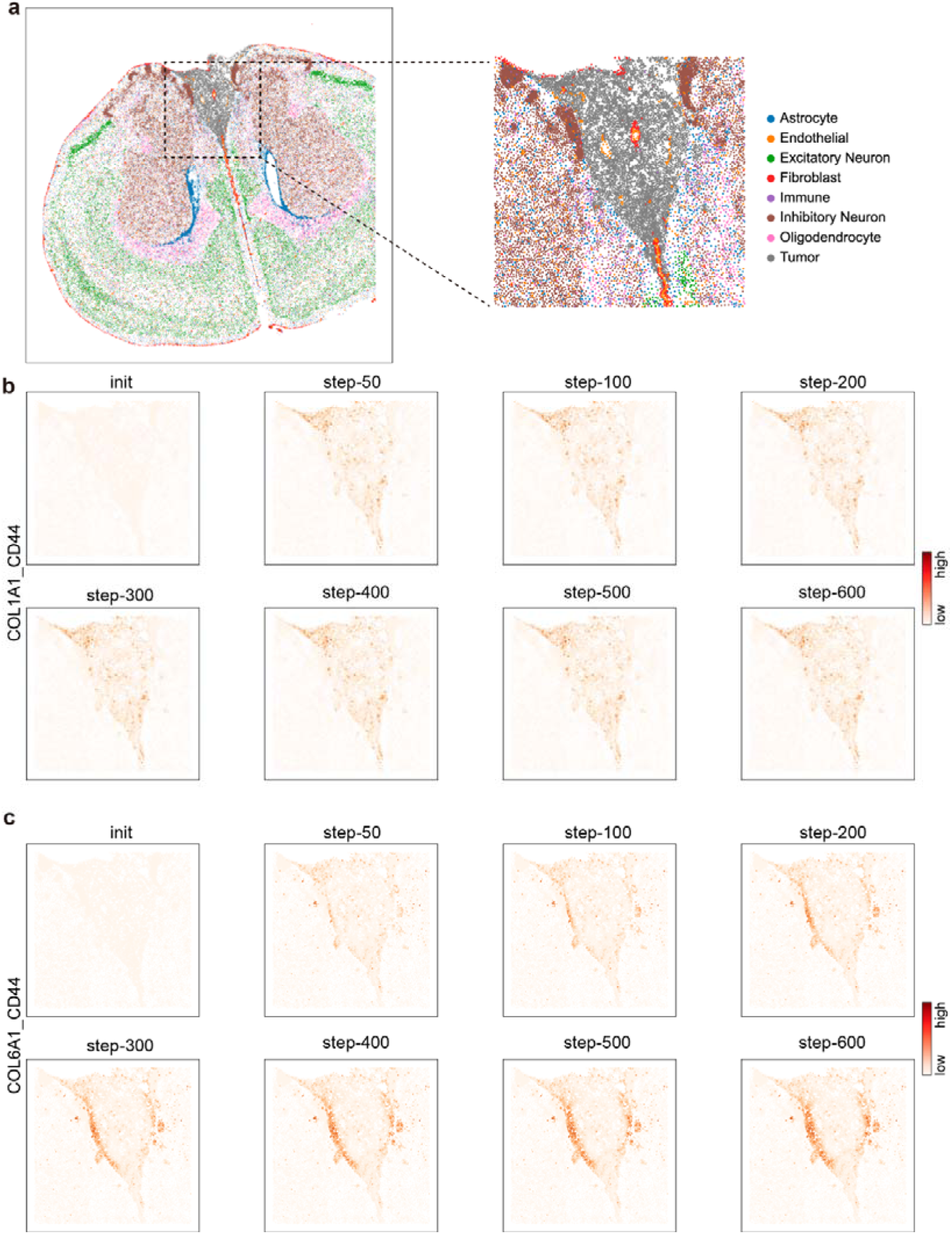
| Cell type annotation and dynamics changes of interactions. **a**, Cell type annotation of the selected region in spatial transcriptomics tissue section. The left is original dataset and the right is selected region containing tumor tissue. **b**, Dynamics visualization of interaction intensities between *Col1a1* and *Cd44* over the SpaGRD simulation process. Each panel represents the interaction map at a specific diffusion-reaction step, showing progressive refinement of spatial interaction estimation. **c**, Same as in **b**, but for the COLA1-CD44 ligand-receptor pair.

**Supplementary Fig. 7.**
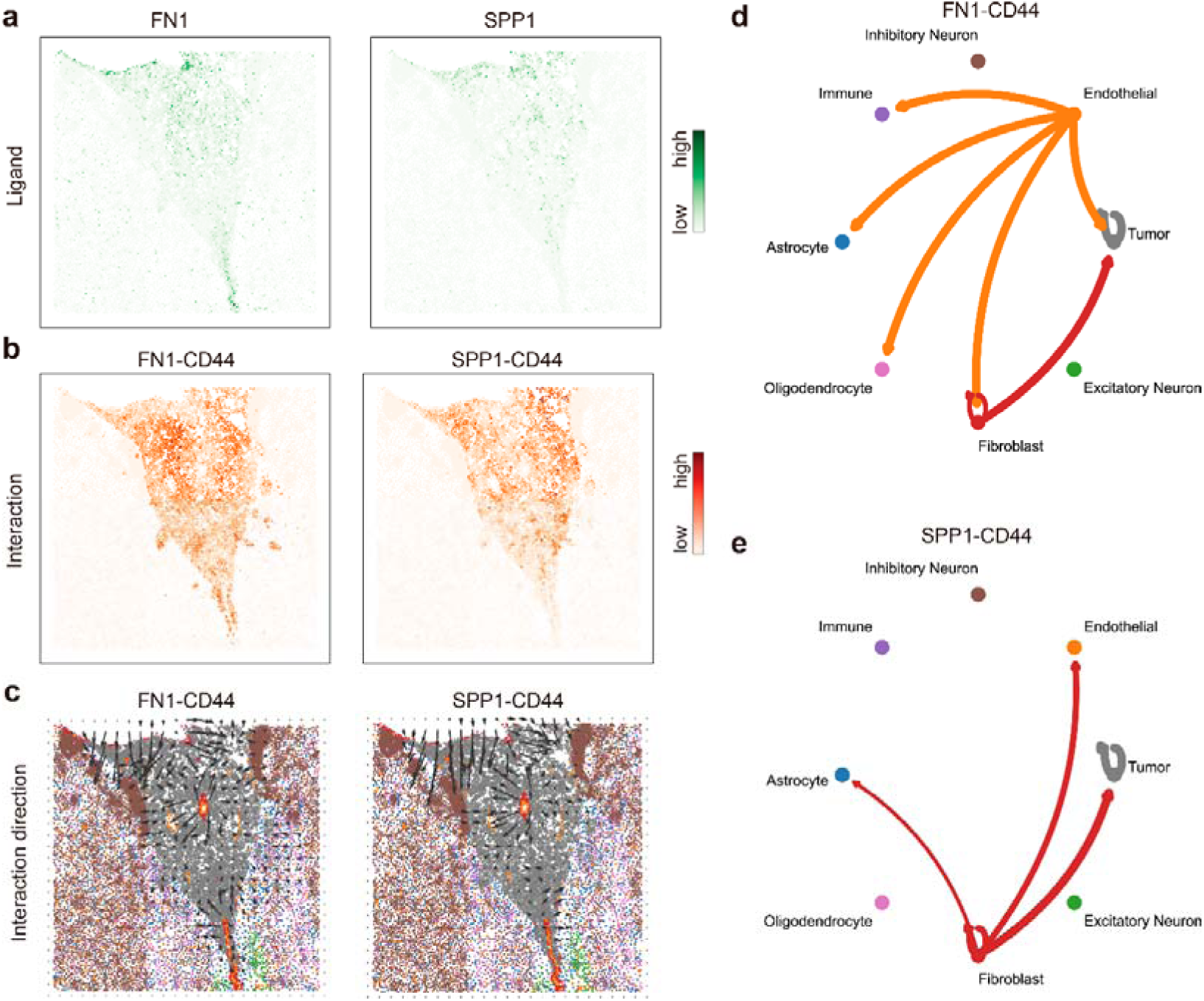
| FN1–CD44 and SPP1–CD44 interaction analysis using SpaGRD. **a**, Spatial expression patterns of the ligands *Fn1* and *Spp1* in the tumor microenvironment, showing localized enrichment around fibroblast and tumor regions. **b**, SpaGRD-inferred interaction intensity maps for the FN1-CD44 and SPP1-CD44 ligand-receptor pairs. Both interactions exhibit strong signals within and around the tumor core. **c**, Vector fields indicating the signaling directions of inferred interactions. **d**, Communication network of FN1-CD44 interactions, with directed edges representing ligand-to-receptor signaling. Fibroblasts act as the major source of signals targeting tumor, endothelial, and excitatory neuron compartments. **e**, Communication network for the SPP1-CD44 pair, highlighting fibroblast-originated signaling primarily targeting other cell types.

**Supplementary Fig. 8.**
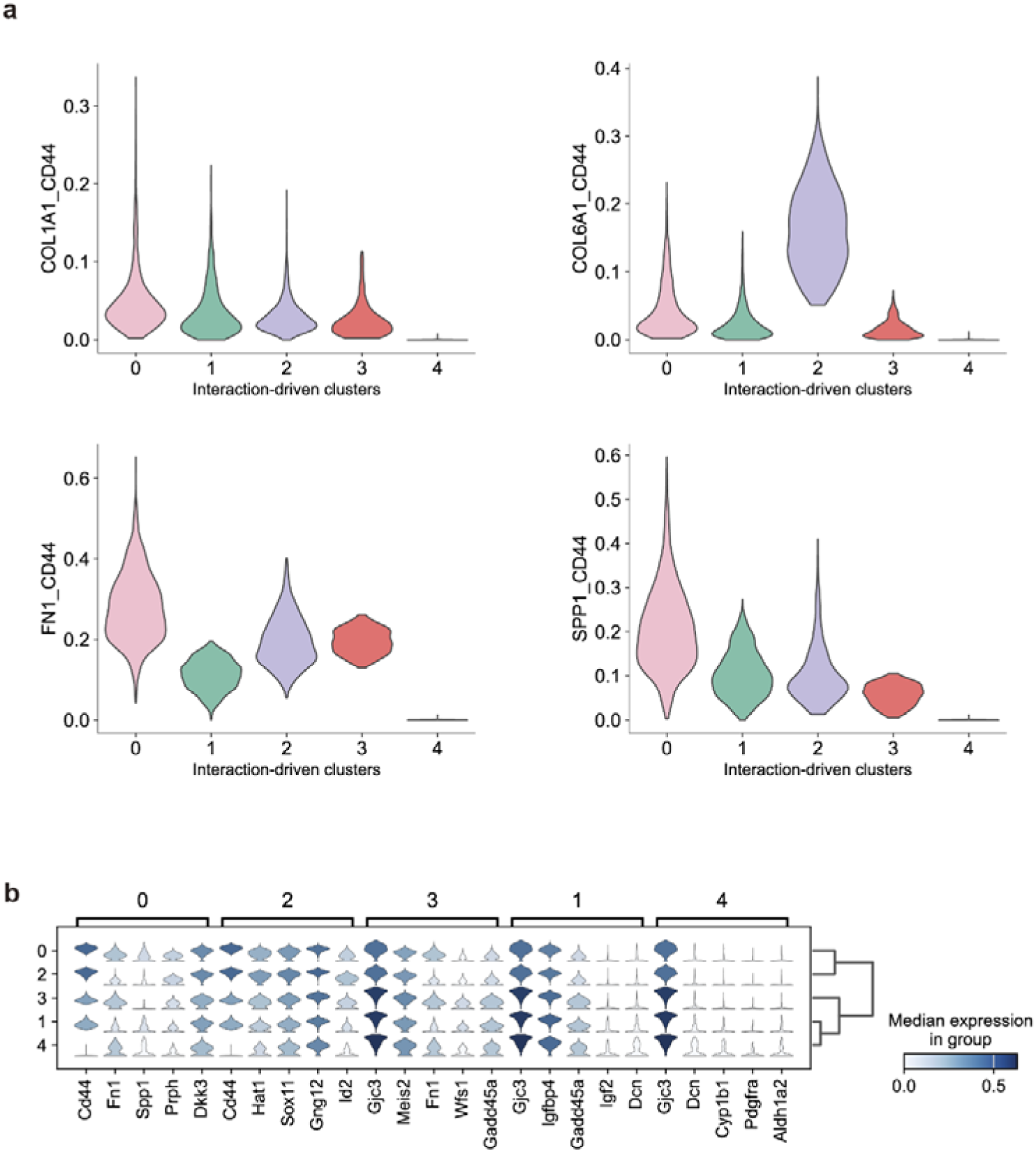
| Interaction and gene expression differences between the interaction-driven tumor clusters. **a**, The violin plots of interaction differences in these clusters. The cluster 0 showed higher COL1A1-CD44 interactions. The cluster 2 showed higher COL6A1-CD44 interactions. The cluster 0 and cluster 2 showed higher FN1-CD44 interactions. The cluster 0 and cluster 1 showed higher SPP1-CD44 interactions. **b**, The violin plot of gene expression differences in these interaction-driven tumor clusters.

**Supplementary Fig. 9.**
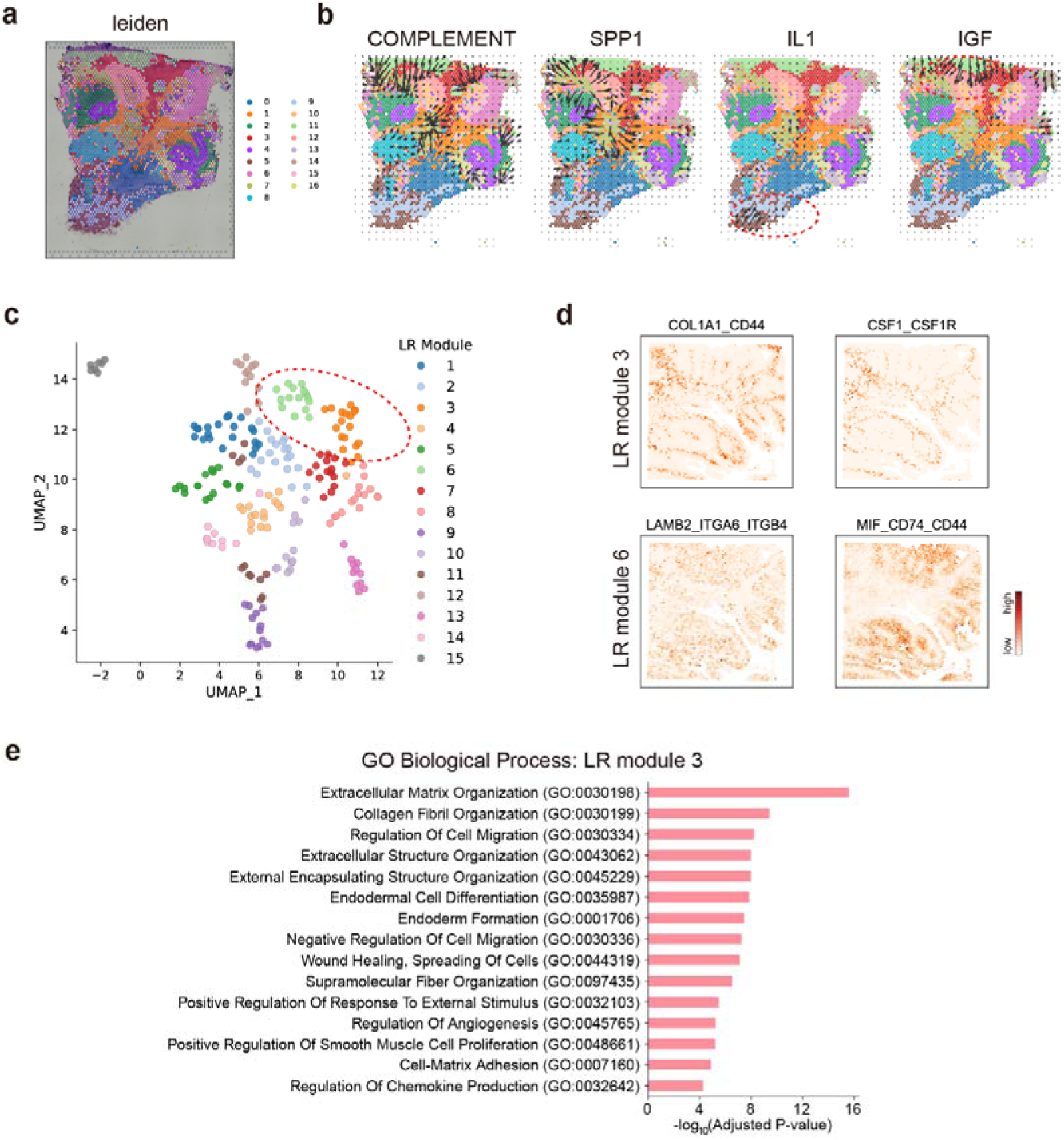
| Communication network and LR module. **a**, Cluster result based on spatially variable gene expression by Leiden. **b**, Communication network of COMPLEMENT (Complement System), SPP1 (Secreted Phosphoprotein 1), IL1 (Interleukin-1), IGF (Insulin-like Growth Factors) interactions, with directed edges representing pathway signaling. **c**, The UMAP visualization of clustering spatially variable ligand-receptor interactions. 15 ligand-receptor (LR) modules were identified. Each point represents a ligand-receptor pair. **d**, Interaction intensity maps inferred by SpaGRD for the ligand-receptor pairs COL1A1-CD44 and CSF1-CSF1R on LR module 3, and LAMB2-ITGA6-ITGB4 and MIF-CD74-CD44 on LR model 6. **e**, Enriched GO terms of genes positively correlated with ligand-receptor pairs in LR module 3.

**Supplementary Fig. 10.**
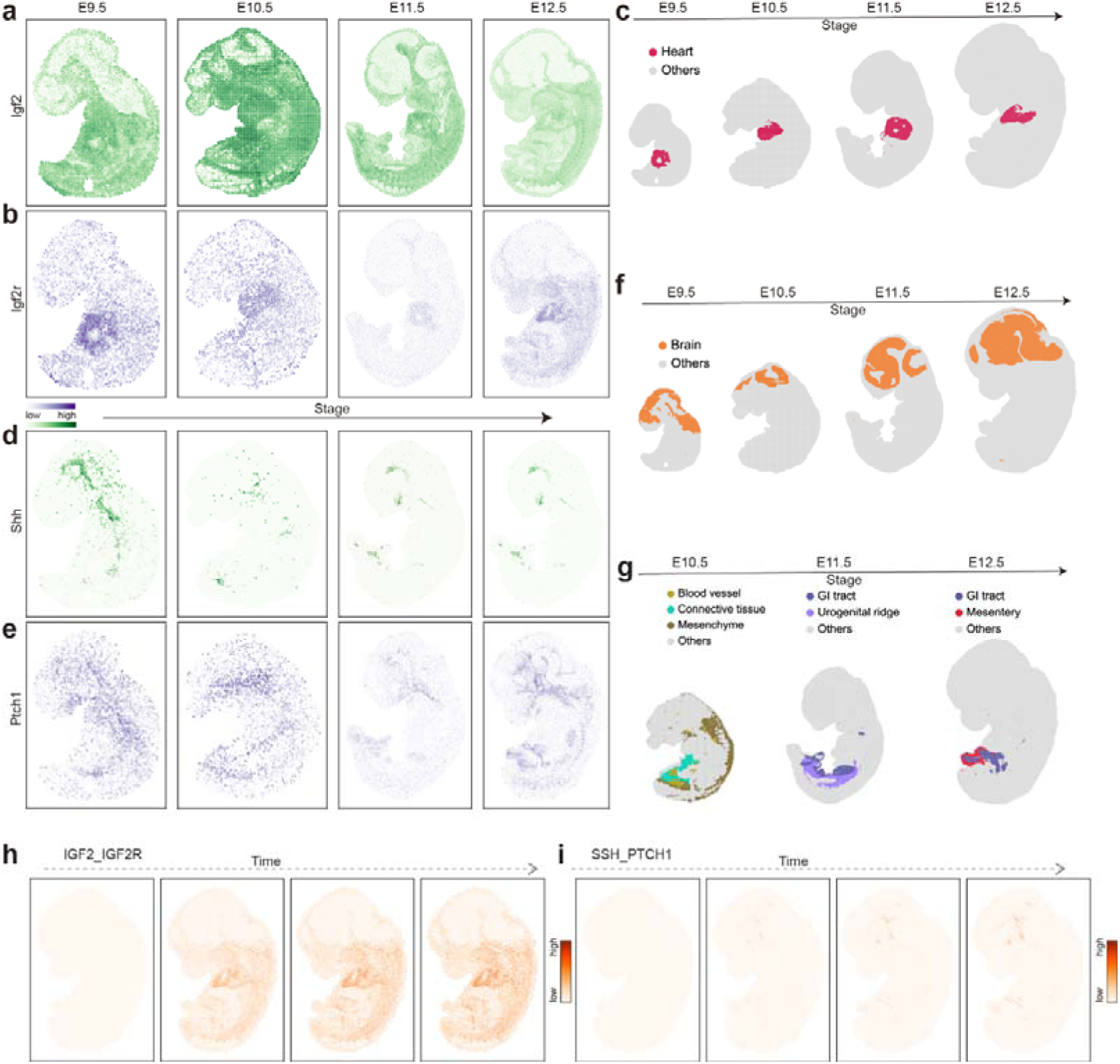
| Spatial expression and interaction patterns of IGF2-IGF2R and SHH-PTCH1 during mouse embryonic development. **a**, Spatial expression pattern of *Igf2* across developmental stages E9.5 to E12.5, showing widespread and dynamic distribution. **b**, Spatial expression pattern of *Igf2r* across stages, with concentrated expression at E9.5 and E10.5, especially in heart regions. **c**, Tissue annotations highlighting heart regions across four stages, which align with regions of high IGF2-IGF2R interaction. **d**, Spatial expression pattern of *Shh*, showing localized expression in the brain and gastrointestinal regions at early stages. **e**, Spatial expression pattern of *Ptch1*, showing spatial overlap with Shh, particularly at E9.5 and E10.5. **f**, Tissue annotations highlighting brain regions across stages, indicating potential areas of active SHH-PTCH1 signaling. **g**, Tissue annotations highlighting gastrointestinal tract, urogenital ridge, mesenchyme, and related tissues that correspond with SHH-PTCH1 co-expression regions. **h**, IGF2-IGF2R interaction intensity maps inferred by SpaGRD from E9.5 to E12.5, showing strong activity in heart and surrounding regions. **i**, SHH–PTCH1 interaction intensity maps inferred by SpaGRD, showing high activity in brain and gastrointestinal regions.

**Supplementary Fig. 11.**
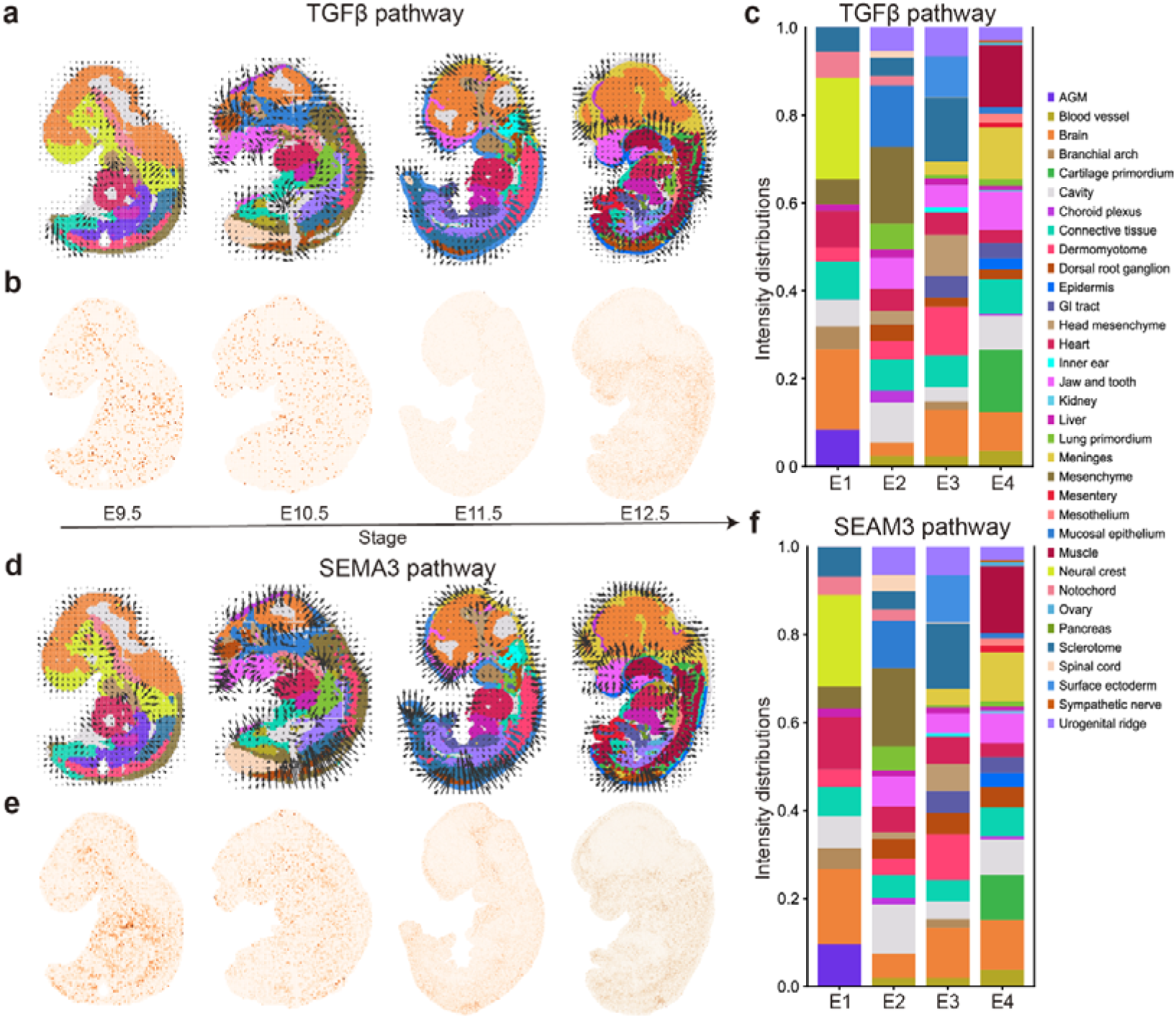
**| Vector field and spatial distributions of TGF**β **pathway and SEMA3 pathway. a**, Vector fields indicating the spatial flow of inferred TGFβ signaling pathway. **b**, Spatial intensities of the TGFβ signaling pathway. **c**, Intensity distribution of TGFβ pathway activities across four developmental stages. **d**, Vector fields indicating the spatial flow of inferred SEMA3 signaling pathway. **e**, Spatial intensities of the SEMA3 signaling pathway. **f**, Intensity distribution of SEMA3 pathway activities across four developmental stages.

**Supplementary Fig. 12.**
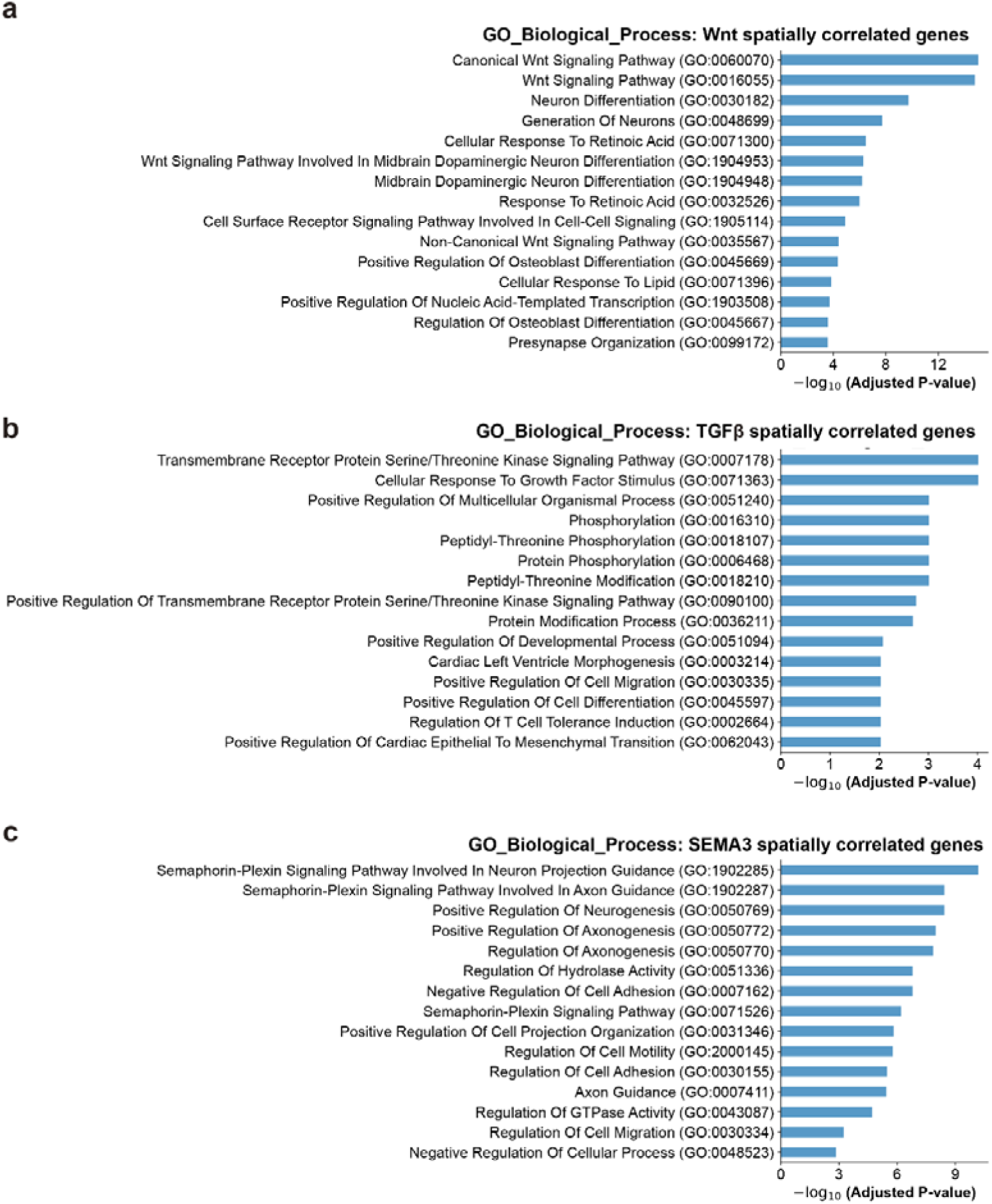
| Gene Ontology enrichment analyses for gene sets that are spatially correlated to signaling pathways. **a**, GO enrichment analysis of genes positively correlated with Wnt signaling pathway. **b**, GO enrichment analysis of genes positively correlated with TGFβ signaling pathway. **c**, GO enrichment analysis of genes positively correlated with SEMA3 signaling pathway.

